# A series of Orai1 gating checkpoints in transmembrane and cytosolic regions requires clearance for CRAC channel opening: Clearance and synergy of Orai1 gating checkpoints controls pore opening

**DOI:** 10.1101/2020.07.16.207183

**Authors:** Adéla Tiffner, Romana Schober, Carmen Höglinger, Daniel Bonhenry, Saurabh Pandey, Victoria Lunz, Matthias Sallinger, Irene Frischauf, Marc Fahrner, Sonja Lindinger, Lena Maltan, Sascha Berlansky, Michael Stadlbauer, Rainer Schindl, Rudiger Ettrich, Christoph Romanin, Isabella Derler

**Author notes:** Corresponding author: (ID).

## Abstract

The initial activation step in gating of ubiquitously expressed Orai1 Calcium (Ca^2+^) ion channels represents the store-dependent coupling to the Ca^2+^ sensor protein STIM1. An array of constitutively active Orai1 mutants gave rise to the hypothesis that STIM1 mediated Orai1 pore opening is accompanied by a global conformational change of all Orai TM helices within the channel complex. Here, we prove that a local conformational change spreads omnidirectionally within the Orai1 complex. Our results demonstrate that a global, opening-permissive allosteric communication of TM helices is indispensable for pore opening and requires clearance of a series of Orai1 gating checkpoints. We discovered these gating checkpoints in middle and cytosolic extended TM domain regions. Our findings are based on a library of double point mutants that contain each one loss-of-function (LoF) with one gain-of-function (GoF) point mutation in a series of possible combinations. We demonstrated that an array of LoF mutations act dominant over most GoF mutations within the same as well as of an adjacent Orai subunit. We further established inter- and intramolecular salt-bridge interactions of Orai subunits as a core element of an opening-permissive Orai channel architecture. Collectively, clearance and synergistic action of all these gating checkpoints is required to allow STIM1 coupling and Orai1 pore opening.

**Graphical Abstract:** 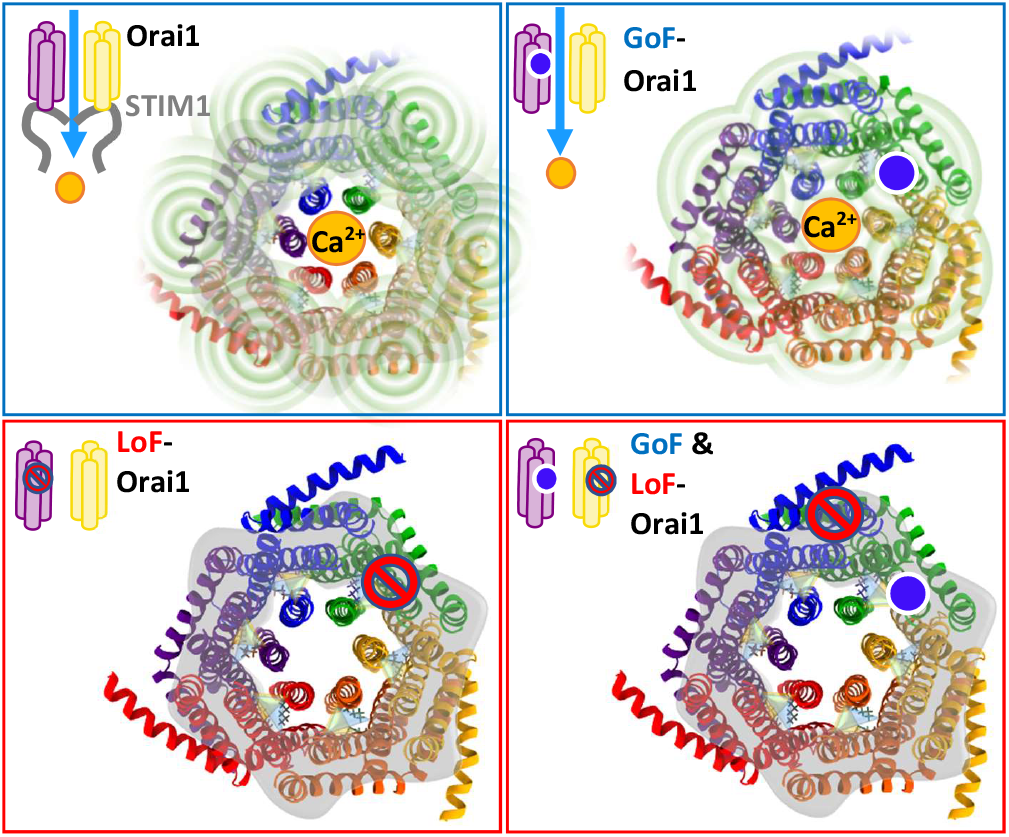

## Introduction

Calcium (Ca^2+^) ions are essential second messengers in the cell that control a variety of processes in immune cells and other cell types^1–3^. One main Ca^2+^ entry pathway into the cell represents the Ca^2+^ release-activated Ca^2+^ (CRAC) channel, which is activated in response to intracellular Ca^2+^ store-depletion^4,5^.

CRAC channels are composed of two molecular key players, the Orai proteins (Orai1 – Orai3), forming the Ca^2+^ ion channel in the plasma membrane and the stromal interaction molecule 1 (STIM1), a Ca^2+^ sensor in the membrane of the endoplasmic reticulum (ER)^5–17^. Upon Ca^2+^ store depletion of the ER, a signaling cascade is initiated finally leading to the coupling of STIM1 oligomers to Orai channels^16,18–24^ which enables Ca^2+^ ions to enter the cell^25,26^.

Each Orai subunit is composed of four transmembrane domains (TM1-TM4) which are flanked by their N- and C-terminal strands and connected via one intracellular (TM2-TM3) and two extracellular (TM1-TM2, TM3-TM4) loops^17,27,28^. All three cytosolic segments are indispensable for STIM1 mediated Orai activation. The Orai C-terminus functions as the main binding site for STIM1 C-terminus^20,25,29–33^. Concerning the N-terminus and the loop2, it is not yet clear whether they form weak STIM1 binding sites or affect STIM1 binding allosterically^32,34^. Currently, we know that the loop2 region is critical in STIM1 mediated Orai1 gating^35^ and it controls the interplay with the Orai N-terminus to adjust a permissive Orai channel conformation^34^. Furthermore, the Orai N-terminus communicates with STIM1 to govern the authentic CRAC channel hallmarks^36^.

The three currently published crystal and one cryo-EM structure(s) of *drosophila melanogaster* Orai (dOrai) determine that Orai ion channels form hexameric assemblies^37–39^. The Ca^2+^ pore is formed by six TM1 domains centered in the middle of the channel complex. They are surrounded by the TM2 and TM3 regions in a second ring and by TM4 in a third ring. Among the four dOrai structures, one is representative of the closed state^37^. The other three published structures are assumed to constitute open channel variants, as they contain one of the two well-known constitutively activating point mutations H206A (in human Orai1 H134A)^39^ and P288L (in human Orai1 P245L)^38^, respectively. Moreover, these structures unveil unique features of the cytosolic strands. The last 20 amino acids of Orai1 N-terminus form a helical extension of TM1, thus contributing to the cytosolic part of the pore. The C-termini form helical regions connected to TM4. Comparison of the structures reveals consistent results on potential conformational changes upon Orai activation within the inner part of the channel complex including TM1-TM3. Structural resolutions indicate that the basic region in TM1 expands by approximately 10 Å from the closed to the open state. On the contrary, the structural changes upon pore opening of TM4 and the C-terminus at the outmost side of the channel complex are still a matter of debate, due to discrepancies in the structural resolutions^37–39^.

The initial signal for Orai pore opening represents STIM1 coupling to Orai1 C-terminus^31,40,41^. This might lead to conformational changes of the C-termini in the Orai complex which are further transmitted to the pore region. Site-directed and cysteine scanning mutagenesis studies exhibited that not only amino acids in the pore-lining TM1, but overall 15 positions within all TM domains contribute to the maintenance of the closed state^31,34,36,42–50^, as their single point mutations can lead to constitutive activity. Critical regions for Orai1 pore opening represent the hinge connecting TM4 and the C-terminus, the TM2/3 ring, a hydrophobic cluster at the TM1-TM2/3 interface, a serine ridge at the TM1-TM2/3 interface, the H134 residue and the N-terminus^36,51,52^. Pore opening is assumed to be facilitated by a rotation of the hydrophobic region in TM1^52^ as well as by the presence of basic residues at the inner pore^46,53^. Finally, signal propagation from the outmost side to the pore in the center of the channel complex has been suggested to involve inter-dependent conformational changes of all TM helices^51^. A recent computational model indicates a “twist-to-open” Orai gating mechanism, which involves collective counterclockwise rotation of the extracellular part and two independent types of motions of the intracellular side of the channel complex^54^.

In our study, we provide solid proof for this inter-dependent communication of the Orai TM helices using a library of Orai double point mutants systematically combining one gain (GoF)-and one loss-of-function (LoF) single point mutation **(Table 1)**. We hereby present a map of critical checkpoints that are all required to adopt an opening-permissive conformation to guarantee pore opening. Our characterization of a variety of LoF mutations distilled certain basic and acidic residues in the cytosolic portions of TM domains in wild-type Orai1 which form triangular salt bridge interactions and possess a pivotal role in pore opening and functional STIM1 coupling.

**Table 1:**
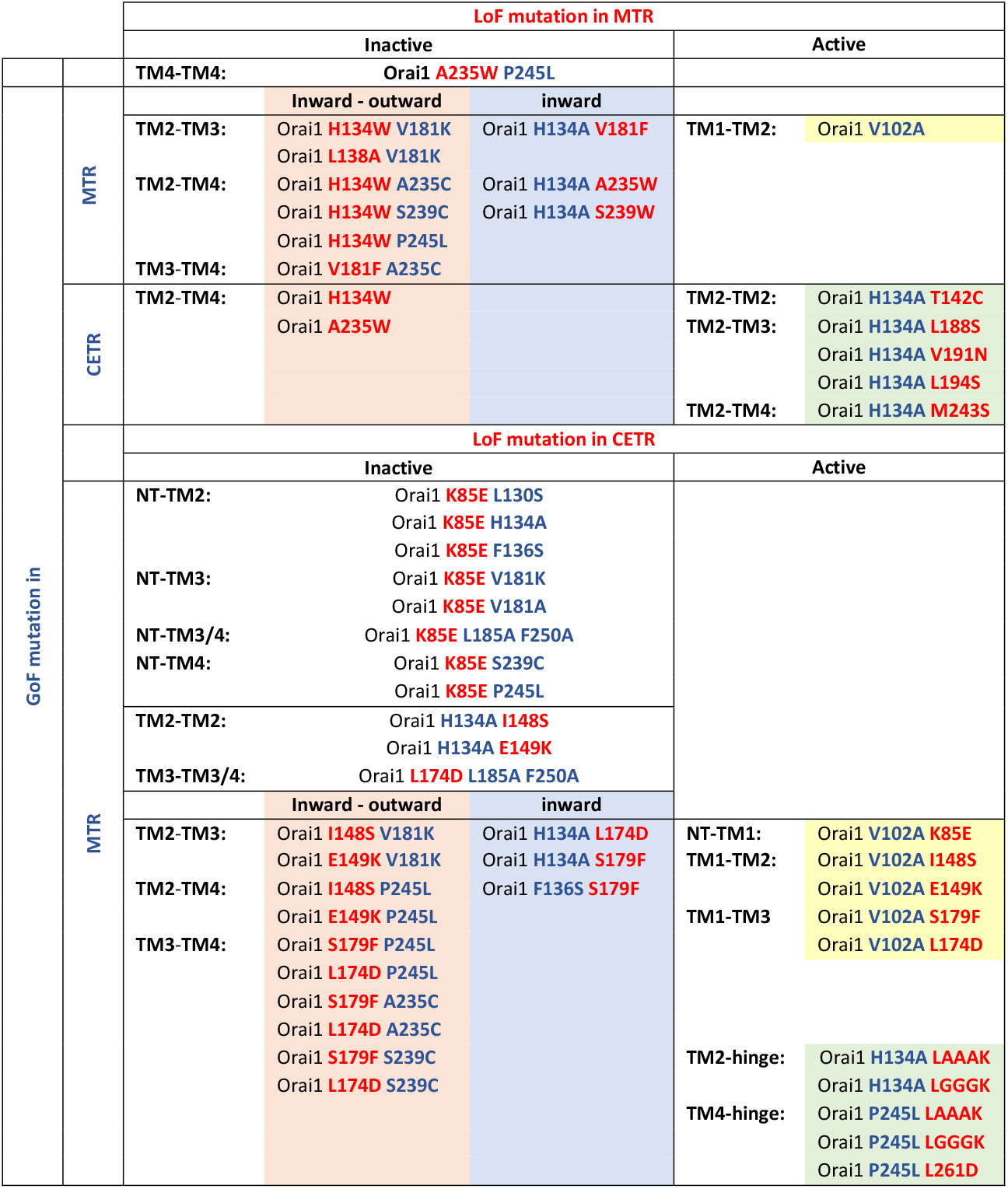
Summary on all tested Orai1 mutants

## Results

### Mutation of several residues located in the middle of TM2, TM3 and TM4 of Orai1 channels can lead either to GoF-or LoF-mutants

Several single point mutants of Orai1, containing amino acid substitutions, in distinct TM regions, have already been reported to display constitutive activity^32,36,41,43,45–47,50–52^. This led to the hypothesis that Orai channel opening is controlled by several critical sites within all four TM regions^35,36,46,51,55,56^. We screened residues in TM2/3 and 4 via single point mutations, with the focus on those located in close proximity (2 – 4 Å) within one or between two adjacent Orai subunits **(Table 2, 3**; include distances estimated via Pymol in the hOrai1 model^46,57^ based on the X-ray structure of the closed dOrai PDB ID: 4HKR). We assumed that they are most likely involved in manifesting a stable closed or open conformation^58^ by forming noncovalent interactions with each other. As the mutation of those sites should not disturb the TM-helix itself, most residues, with a few exceptions (e.g. V181K, S239W), were exchanged by ones with a preference for helix formation such as serine or alanine, but not by beta-branched residues which might disrupt the helix^59^. We discovered, among residues located in close proximity of adjacent TM domains **(Table 2, 3)**, an overall number of 13 positions in the Orai1 helices TM2, TM3 and TM4 that lead to constitutive activity upon single point mutation **(Fig 1a, Supp Fig 1 ac)**. Our screen is largely in line with the cysteine screen of Yeung et al.^51^ except for some variations. We elucidated the following, novel gain-of-function (GoF) single point mutations within Orai1: in TM2 L130S, F136S, in TM3 W176S, V181S/K, F187S **(Fig 1 a, b, c; Supp Fig 2I a-j; Supp Fig 2II a-j)**, in addition to the reported H134A, A137V^46^, L138F^47^, W176C^48^, V181A, L185A^34,36^ and diverse cysteine substitutions^51^. The constitutive TM4 mutants Orai1 A235C, S239C, F250C^51^ and P245L^41^ behaved in an identical manner to the recently reported ones **(Fig 1 a, e, f, h, i; Supp Fig 2I k-r; Supp Fig 2II k-r)**. In general, except for the positions A137 and L138, a substitution of a strongly hydrophobic residue to a serine, alanine or cysteine destabilizes the closed Orai1 channel conformation and induces constitutive activity.

**Figure 1:**
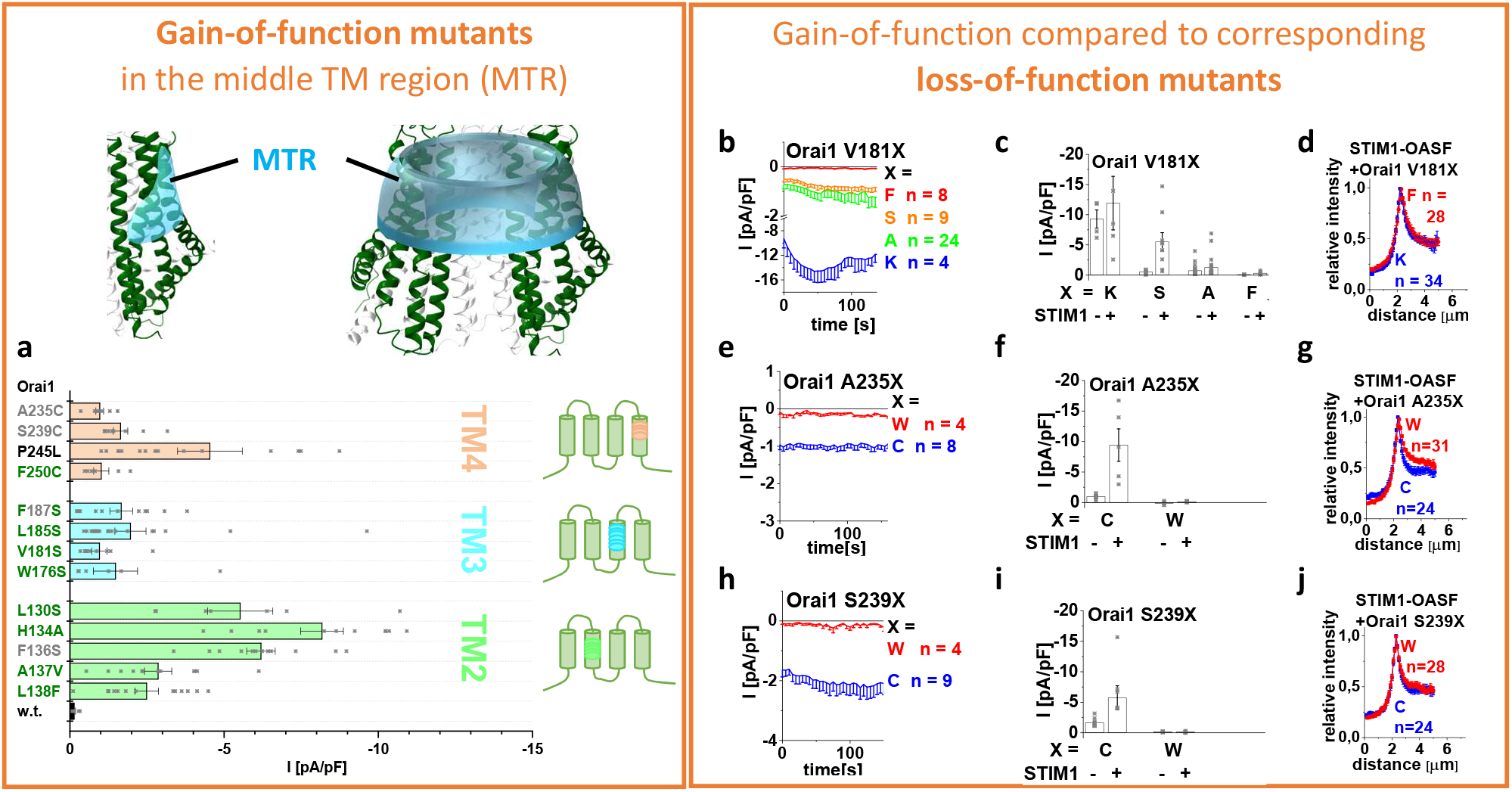
A screen on Orai1 TM domains revealed several GoF mutations and for some the corresponding LoF mutations. Scheme representing the location of the MTR region (cyan) within one Orai subunit (left) and the whole channel complex (right). a) Block diagram showing current densities of constitutively active point mutants in TM2 (green), TM3 (blue) and TM4 (orange) *(n* = 6 – 19 cells; values are mean ± SEM). Current densities of constitutive GoF mutant are significantly different to those of Orai1 (p < 0,05). The colour code used for the mutations refers to their close proximity to a residue within one subunit (green) or an adjacent subunit (gray). P245L is shown in black as it causes constitutive activity, but is not located in close proximity to residues of adjacent TM domains. b) e) h) Time course representing current densities of Orai proteins containing the LoF-mutation V181F in comparison to the GoF-mutations V181S, V181A, V181K (b), the LoF-mutation A235W in comparison to the GoF-mutation A235C (e) and the LoF-mutation S239W in comparison to the GoF-mutation S239C (h) in the absence of STIM1. c) f) i) Block diagram with current densities of mutants from (b), (e) and (h) in the absence (t = 0 s) compared to the presence (maximum current density) of STIM1. Currents of GoF mutants are significantly different to those of the corresponding LoF mutants (p < 0,05). d) g) j) Intensity plots of STIM1-OASF co-expressed with Orai1 V181K compared to Orai1 V181F (d), with Orai1 A235C compared to Orai1 A235W (g) and with Orai1 S239C compared to Orai1 S239W (j) (at 4 μm, p > 0,05).

**Table 2:**
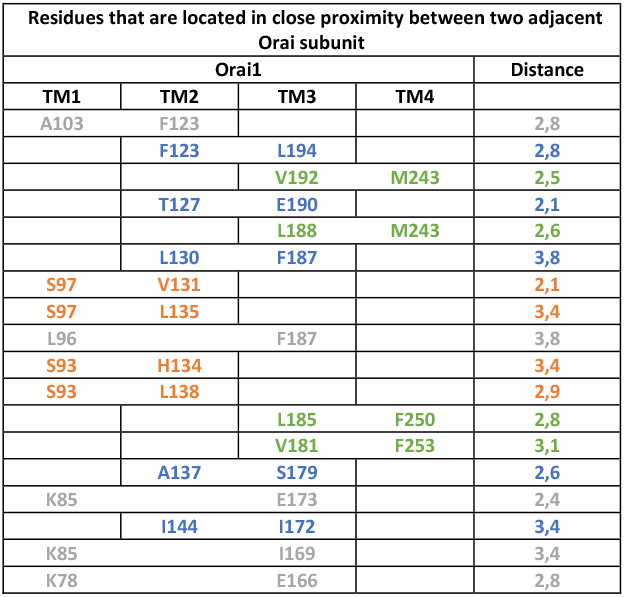
Residues that are located in close proximity within one Orai subunit. Color code: Pairs of residues located in close proximity between TM1-TM2 (orange), TM1-TM3 (gray), TM2-TM3 (blue) and TM3-TM4 (green) and corresponding distance of the respective side chains.

In contrast, the exchange of a few of those residues by ones with strongly distinct properties of size and/or hydrophobicity can lead to loss of function (LoF) mutants **(Supp Fig 2I a-r; Supp Fig 2II a-r)**. This fact has already been reported for Orai1 H134A and Orai1 L138F, which are constitutively active, while Orai1 H134W and L138A remain inactive also in the presence of STIM1^46^. Moreover, we discovered loss of function of Orai1 V181F in TM3, Orai1 A235W and S239W in TM4, both, in the absence **(Fig 1 b, c, e, f, h, i; Supp Fig 2 I&II e, k, m)** as well as the presence of STIM1 **(Fig 1 c, f, i; Supp Fig 2 I&II f, l, n)**. It seems that especially large, bulky residues can lead to a loss of function, however, due to exceptions (L138F, **Fig 1a, Supp Fig 2 II**) no clear rules can be postulated. There is no obvious dependence of gain- or loss-of-function in terms of hydrophobicity (Kyte and Doolittle scale^60^) of the introduced residue in line with previous findings (**Supp Fig 2 I)**^58^. At some other positions, such as L185, F187 and P245, any amino acid substitution leaves the Orai1 channel in a functional state, either constitutively or activatable by STIM1 **(Supp Fig 2 I&II g-j, o, p)**^41^. Also, the substitution of L130 and F136 by the small polar serine or the bulky, hydrophobic tryptophan retained both store-operated activity **(Supp Fig 2 I&II a-d)**.

To investigate whether loss of function of above-mentioned mutants is partially a result of impaired STIM1 coupling, we investigated the intensities of the **O**rai1 **a**ctivating **S**TIM1 C-terminal **f**ragment, OASF (aa 233 – 474), in the presence of diverse LoF compared to GoF mutants. However, both, GoF and corresponding LoF point mutants revealed comparable co-localization intensities with OASF **(Fig 1 d, g, j)** compatible with similar affinities among the Orai1 mutants.

In summary, our screening on nearby residues from adjacent TM domains within and between Orai1 subunits led to the discovery of in total 13 positions in Orai1 TM2/3/4, that contribute to the establishment of the closed state of the Orai1 channel. They can become constitutively active upon single point mutation to certain amino acids independent of the presence of STIM1. Some of them are additionally involved in the maintenance of an opening-permissive pore geometry or signal propagation, since distinct amino acid substitutions at those positions can lead to loss of function, without interfering with the coupling to STIM1-OASF. Overall, all these positions are concentrated in a conical ring spanning across the middle plane of the TM2/3/4 domains and a segment of TM4 closer to the extracellular side, which we call middle transmembrane region (MTR) **(Fig 1, scheme of subunit and channel).**

### MTR-LoF Orai1 point mutations act dominant over MTR-GoF mutations independent of their location relative to each other

Orai1 channel activation is initiated via STIM1 coupling to the Orai1 C-terminus^49^, however, how this activation signal is transmitted to the pore has so far remained unclear. The huge variety of constitutively active Orai1-TM-mutants **(Fig 1, Supp Fig 1, Supp Fig 2 I&II)**^36,41,45–47,50–52^ allows to assume that STIM1 binding successively alters the conformations of TM4 up to TM1 via interdependent TM motions, finally leading to pore opening. Here, we exploited the various MTR-GoF and MTR-LoF point mutations to determine how a GoF mutation-initiated activation signal propagates through the channel complex to establish a pore opening. Therefore, we generated and investigated a pool of double point mutants including one MTR-GoF and one MTR-LoF point mutation in distinct TM domains, one of which is located in a TM region (e.g. TM2) closer to the pore than the other one (e.g. TM4) **(Table 1)**. Initially, we focused on the positions H134 in TM2 and S239 in TM4. As anticipated, the MTR-LoF point mutation H134W in TM2 combined with the MTR-GoF point mutation S239C in TM4 (Orai1 H134W S239C) led to the loss of function in the absence of STIM1 **(Fig 2 a, b)**. Further, the coexpression of STIM1 was unable to restore the function **(Fig 2 b)**. Analogously, other combinations with the LoF point mutation closer to the pore, like Orai1 H134W V181K, Orai1 L138A V181K, Orai1 H134W A235C, Orai1 H134W P245L and Orai1 V181F A235C **(Fig 2 e)**, abolished or significantly reduced STIM1 mediated activation. Interestingly, the double point mutant including the MTR-GoF point mutation H134A in TM2 combined with the MTR-LoF point mutation S239W in TM4 remained also inactive independent of the presence of STIM1 **(Fig 2 f, g)**. Similar combinations with the MTR-GoF point mutation closer to the pore such as Orai1 H134A V181F, Orai1 H134A A235W/S239W showed significantly reduced or eliminated function also in the presence of STIM1 **(Fig 2 j)**. Moreover, we tested a double point mutant containing the MTR-GoF and the MTR-LoF point mutation within the same TM domain, specifically TM4. Orai1 A235W P245L also displayed loss of constitutive activity **(Supp Fig 3 i)**, indicating that the diverse MTR-LoF point mutations act in a dominant manner. Control experiments revealed that these non-functional double point mutants displayed comparable plasma membrane expression like the associated constitutive single point mutants **(Supp Fig 3 a, e, h, k)**. Additionally to all MTR-GoF mutants, one GoF mutant containing substitutions within the hinge region, Orai1 ANSGA (L261A-V262N-H264G-K265A) is currently known^49^. Its constitutive activity can also be overruled by MTR-LoF point mutations (Orai1 H134W ANSGA; Orai1 A235W ANSGA; **Supp Fig 4 n**). Collectively, MTR-LoF mutations overrule the effect of MTR-GoF mutations independent of their position relative to each other, either located in the middle or outer transmembrane ring.

**Figure 2:**
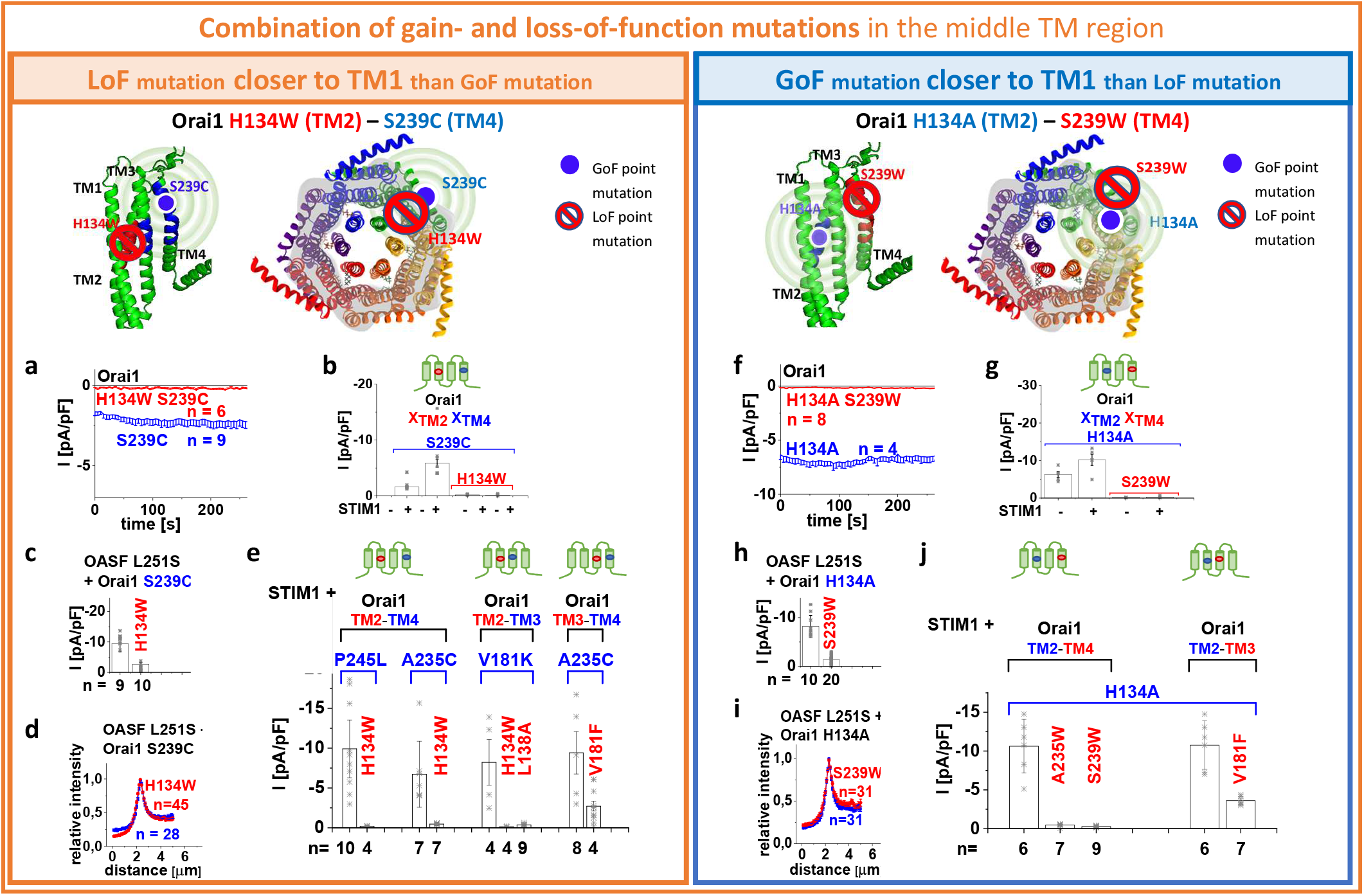
The combination of a GoF and a LoF point mutations in the MTR leads to loss of function of the Orai channel independent of their position relative to each other. Schemes representing the location of the investigated residues within a single subunit (top – left) of Orai1 or the whole channel complex (top – middle), for either LoF or GoF mutation closer to the pore, respectively. Red stop sign represents the position of the LoF mutation, while blue circle shows the position of the GoF mutation. The spheres indicate the impact of the GoF mutation on the entire subunit. In the following, it is valid for (a), (b), (c), (e), (f), (g), (h) and (j) that current densities of MTR-GoF-CETR-LoF double mutants are significantly different of those of the corresponding MTR-GoF mutants (p < 0,05). a) Time course of whole cell current densities of Orai1 S239C compared to Orai1 H134W S239C in the absence of STIM1. b) Block diagram of whole cell current densities of Orai1 S239C, Orai1 H134W S239C in the absence (t = 0 s) and the presence (maximum current densities) of STIM1 (n = 6 – 9 cells; values are mean ± SEM). c) Block diagram of whole cell current densities of Orai1 S239C compared to Orai1 H134W S239C in the presence of STIM1 OASF L251S. d) Intensity plots of STIM1-OASF-L251S co-expressed with Orai1 S239C compared to Orai1 H134W S239C (at 4 μm, p > 0,05). e) Block diagram of whole cell current densities of Orai1 P245L compared to Orai1 H134W P245L; Orai1 A235C compared to Orai1 H134W A235C; Orai1 V181K compared to Orai1 H134W V181K and Orai1 L138A V181K; Orai1 A235C in comparison to Orai1 V181F A235C, all in the presence of STIM1. f) Time course of whole cell current densities of Orai1 H134A compared to Orai1 H134A S239W in the absence of STIM1. g) Block diagram of whole cell current densities of Orai1 H134A, Orai1 H134A S239W in the absence (t = 0 s) and the presence (maximum current densities) of STIM1 (n = 4 – 9 cells; values are mean ± SEM). h) Block diagram of whole cell current densities of Orai1 H134A compared to Orai1 H134A S239W in the presence of STIM1 OASF L251S. i) Intensity plots of STIM1-OASF-L251S co-expressed with Orai1 H134A compared to Orai1 H134A S239W (at 4 μm, p > 0,05). j) Block diagram of whole cell currents of Orai1 H134A compared to Orai1 H134A A235W, Orai1 H134A S239W and Orai1 H134A V181F, all coexpressed with STIM1.

Furthermore, we used the STIM1 C-terminal fragment, STIM1 OASF L251S already exhibiting an open conformation and thus an enhanced affinity to Orai1^26^, to investigate for potential recovery of the function of the inactive Orai1 double mutants. Both, Orai1 H134W S239C and Orai1 H134A S239W revealed in the presence of OASF L251S marginal, but significantly reduced currents compared to the constitutively active, single point mutants Orai1 S239C and Orai1 H134A, respectively **(Fig 2 c, h)**. Colocalization studies revealed almost unaffected coupling of OASF L251S and STIM1 OASF to the two Orai1 double mutants, Orai1 H134W S239C and Orai1 H134A S239W, compared to the constitutively active single point mutants Orai1 S239C and Orai1 H134A, respectively **(Fig 2 d, i; Supp Fig 3 d, g)**. Also, other double point mutants exhibit affinity for STIM1 OASF to a comparable extent as their corresponding constitutively active point mutants **(Supp Fig 3 b, c, f, j)**.

In contrast to the above mentioned GoF mutations, the constitutively active Orai1 TM1-V102A mutant represents an exception. Its function cannot be overruled by LoF mutations in the MTR, as exemplarily tested for Orai1 V102A H134W **(Supp Fig 4 a)**. It shows non-selective currents with a reversal potential (V_rev_) in the range of ~+ 20 mV, comparable to Orai1 V102A currents^32,43^. The co-expression of STIM1 enhanced V_rev_ to ~+ 50 mV, similar to STIM1 mediated Orai1 V102A currents **(Supp Fig 4 b)**^32,43^. In contrast to Orai1 V102A, the constitutive activity of other TM1 mutants Orai1 F99M and Orai1 V107M are overruled by LoF mutations, as proven by Orai1 F99M H134W and Orai1 V107M H134W double mutants, which both exhibit loss of function **(Supp Fig 4 i, j)**.

In summary, our library of GoF-LoF mutants revealed a dominant role of the LoF over the GoF point mutations within the MTR, except the GoF mutation, V102A. This proves, for the first time, that Orai1 activation is driven by an inter-dependent communication of the TM helices and requires a series of Orai1 pore opening-permissive checkpoints in the MTR.

### Novel Orai1 LoF point mutants in TM2, TM3 and TM4

Our site-directed mutagenesis screen through a series of TM-residues located in close proximity to those of adjacent TM domains revealed besides above described MTR-LoF mutations, additional LoF mutations throughout all TM domains **(Table 2, 3; Supp Fig 1 d-f)**. Newly-discovered LoF mutants represent Orai1 T142A/C, I148S (TM2), E149K (TM2), S179F/M/W/T/R/D (TM3), L188S (TM3), L194S (TM3), M243S (TM4) and 3xA (V262A-S263A-H264A), 3xG ((V262G-S263G-H264G)) (hinge region) **(Fig 3 a, b; Supp Fig 4 k; Supp Fig 5 a; Supp Fig 6 b, c, d)** besides the already known LoF mutations L81A, K85E (TM1)^49,61^, E149A/R (TM2)^54,62^, L174D^49^ and V191N^51^ (TM3) and L261D^49^ (hinge region). Their plasma membrane expression remained unaffected as exemplarily tested for the mutants shown in **Supp Fig 6 a**. In the following we investigated the impact of these single point mutations firstly, on the coupling to STIM1-OASF and secondly, on their potential dominance over the MTR-GoF single point mutation, H134A. Among LoF mutations discovered in **Fig 3 a** only those located in the cytosolic, extended TM regions (CETR) **(scheme, green ring)** interfere with both, STIM1 coupling **(Fig 3 c)** and the constitutive activity induced by the MTR-GoF H134A **(Fig 3 d)**. Conversely, other LoF single point mutations elucidated in **Fig 3 a** leave the coupling to STIM1 OASF and the constitutive activity of Orai1 H134A unimpaired **(Supp Fig 5 b – d)**. Additional hinge mutations described above abolish the co-localization with STIM1 **(Supp Fig 4 l)**, but leave the constitutive activity of Orai1 H134A unaffected **(Supp Fig 4 k, m)**.

**Figure 3:**
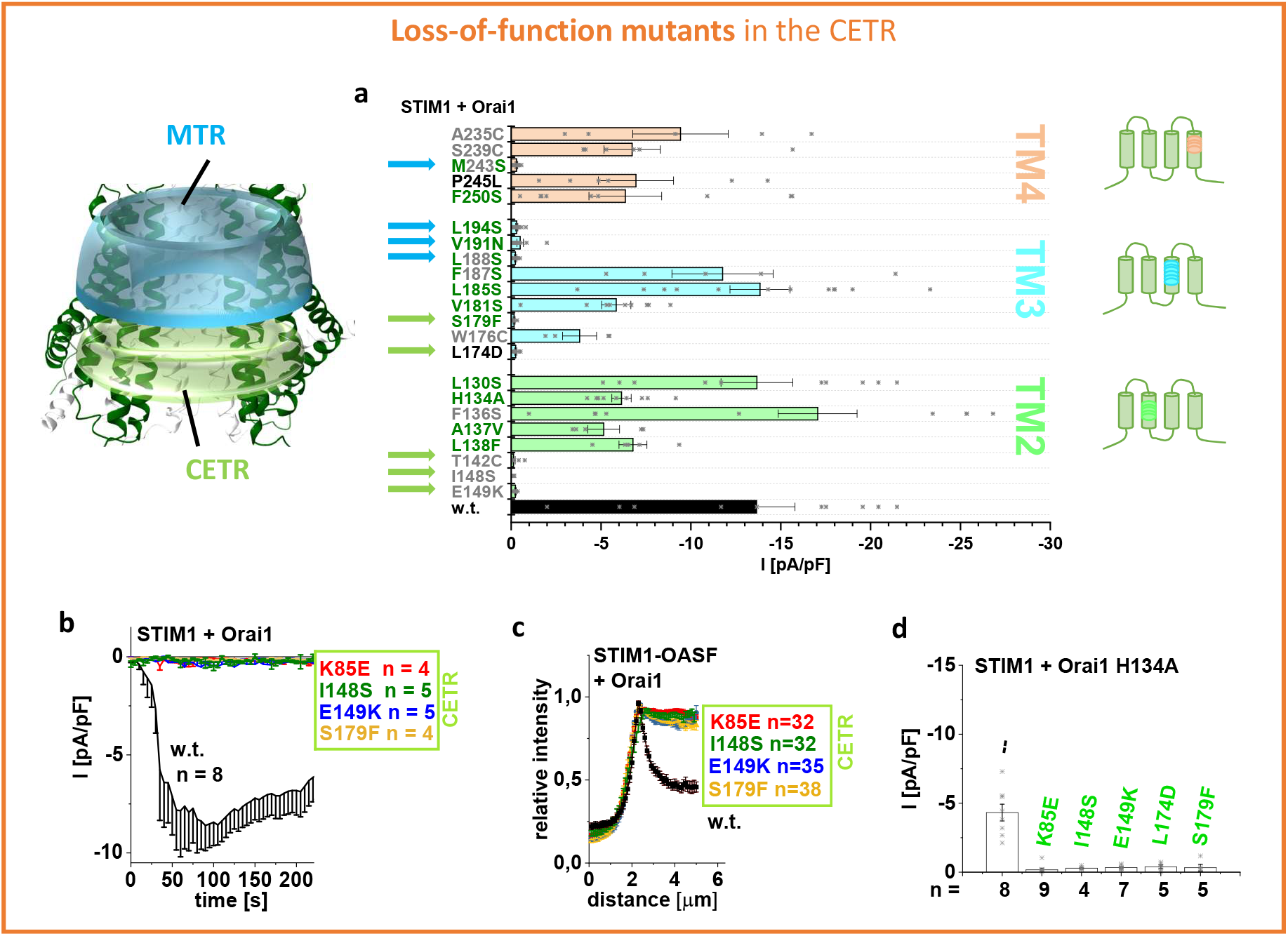
CETR-LoF mutations impair STIM1 coupling and abolish constitutive activity of Orai1 H134A. Scheme representing the location of the MTR region (cyan) and the CETR (green) in the whole channel complex. a) Block diagram showing STIM1 mediated current densities of TM2 (green), TM3 (blue) and TM4 (orange) Orai1 single point mutants (*n* = 4 – 12 cells; values are mean ± SEM). Orai1 T142C, Orai1 I148S, Orai1 E149K in TM2, Orai1 S179F, Orai1 L174D, Orai1 L188S, Orai1 V191N, Orai1 L194S in TM3 and Orai1 M243S represent the critical LoF point mutants (blue arrows point to MTR and green to CETR). The colour code used for the mutants refers to their close proximity to a residue within one subunit (green) or an adjacent subunit (gray). b) Time course of STIM1 mediated CETR LoF Orai1 mutant current densities discovered in (a). STIM1 mediated current densities of the LoF mutants are significantly different compared to those of Orai1 wild-type. c) Intensity plots of STIM1-OASF coexpressed with Orai1 mutants shown in (b) compared to wild-type Orai1 (at 4 μM, p < 0,05 for CETR mutants). d) Block diagram of STIM1 mediated Orai1 double mutant current densities of Orai1 Orai1 K85E H134A, Orai1 H134A I148S, Orai1 H134A E149K, Orai1 H134A L174D and Orai1 H134A S179F in comparison to Orai1 H134A (p < 0,05 for GoF-Orai1-H134A mutants containing I148S, E149K, L174D or S179F).

**Table 3:**
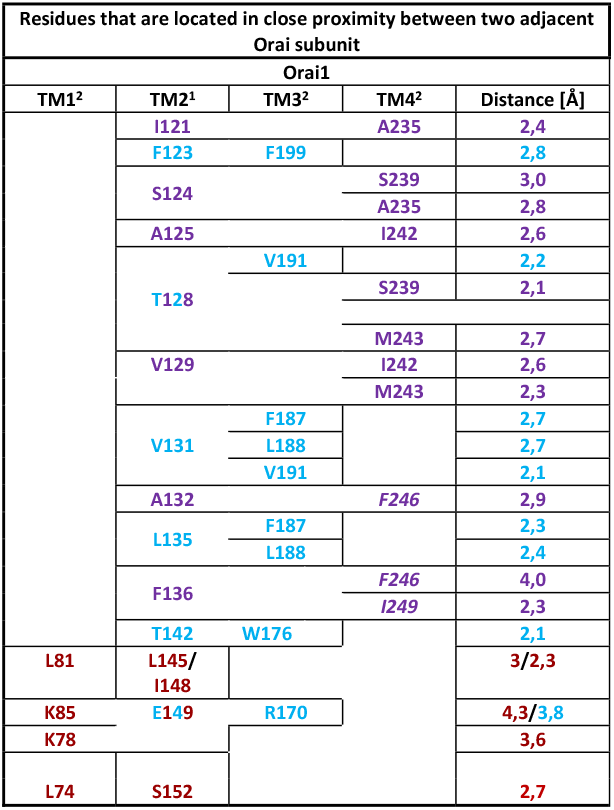
Residues that are located in close proximity between two adjacent Orai subunits. Color code: Pairs of residues located in close proximity between TM1^2^-TM2^1^ (dark red), TM2^1^-TM3^2^ (light blue) and TM2^1^-TM4^2^ (purple) and corresponding distance of the respective side chains.^1^ and ^2^ indicate adjacent Orai1 subunits.

Overall, CETR-LoF mutations possess a multidimensional role including inhibition of STIM1 binding as well as interference with an opening-permissive channel formation and function. Thus, we used these CETR-LoF mutations in the following to investigate their impact on different MTR-GoF mutations.

### N-terminal LoF point mutation K85E abolishes the constitutive activity of all GoF TM2/3/4 mutants

Initially, we investigated the impact of a defect N-terminus on MTR-GoF point mutations employing the prominent N-terminal K85E point mutation leading to loss-of-function of Orai1^61^. We introduced K85E in either a constitutive TM2-, TM3- or TM4-point mutant. All double/triple mutants: Orai1 K85E L130S, Orai1 K85E H134A, Orai1 K85E F136S (in TM2), Orai1 K85E L185A F250A, Orai1 K85E V181K and Orai1 K85E V181A (in TM3/TM4) and Orai1 K85E P245L, Orai1 K85E S239C (in TM4) **(Supp Fig 7 a – c)** displayed loss of function also in the presence of STIM1, in line with Yeung et al.^51^. Plasma membrane expression remained unaffected for all these inactive double/triple mutants **(Supp Fig 7 d)**. Coupling to the STIM1 C-terminal fragment OASF was partially reduced **(Supp Fig 7 e – h)**, which, however, does not account for a total loss of function of the double/triple point mutants. In total, K85E is dominant over the GoF single point mutations and affects functional pore conformation and signal propagation already prior to STIM1 binding.

### CETR-LoF point mutations act dominant over MTR-GoF mutations independent of their location relative to each other

Next, we employed Orai1 CETR-LoF mutations of TM2 and TM3 to investigate their impact on the above described Orai1 GoF mutations located in the MTR. Therefore, we combined LoF and GoF point mutations in distinct TM domains, whereas one was located closer to the pore (e.g. TM2) than the other one (e.g. TM3), with special focus on H134, S239 in the MTR and L174 in the CETR. As expected, the double mutant Orai1 L174D S239C, containing the CETR-LoF-L174D in TM3 closer to the pore than the MTR-GoF-S239C in TM4, exhibited loss of function, both in the absence and presence of STIM1 **(Fig 4 a, b)**. Analogously, other constitutive mutants with a CETR-LoF point mutation located closer to TM1 than the MTR-GoF substitution (Orai1 I148S V181K, Orai1 E149K V181K, Orai1 I148S/E149K P245L, Orai1 L174D/S179F P245L, Orai1 L174D A235C; **Fig 4 e**) displayed also loss of function. Interestingly, vice versa also double mutants with the MTR-GoF point mutation closer to TM1 (e.g. H134A in TM2) than the CETR-LoF point mutation (e.g. L174D in TM3), such as Orai1 H134A L174D, and others (Orai1 H134A S179F, Orai1 F136S S179F) displayed abolished constitutive activity independent of the presence of STIM1 **(Fig 4 f, g, j)**. Moreover, we combined CETR-LoF and MTR-GoF point mutations of the two membrane planes within one TM domain (Orai1 H134A I148S/E149K in TM2, Orai1 L174D L185A F250A in TM3). These mutants also failed to show constitutive activity **(Supp Fig 8 k, n)**. All nonfunctional double mutants displayed comparable plasma membrane localization like the constitutive single point mutant **(Supp Fig 8 a, f, j, m)**. Also, the constitutive activity of the Orai1 ANSGA mutant can be overruled by LoF point mutations in the CETR, similar to the loss of function of Orai1 K85E ANSGA^49^. Collectively, similar to MTR-LoF in Fig 2, also CETR-LoF mutations act dominant over MTR-GoF mutations.

**Figure 4:**
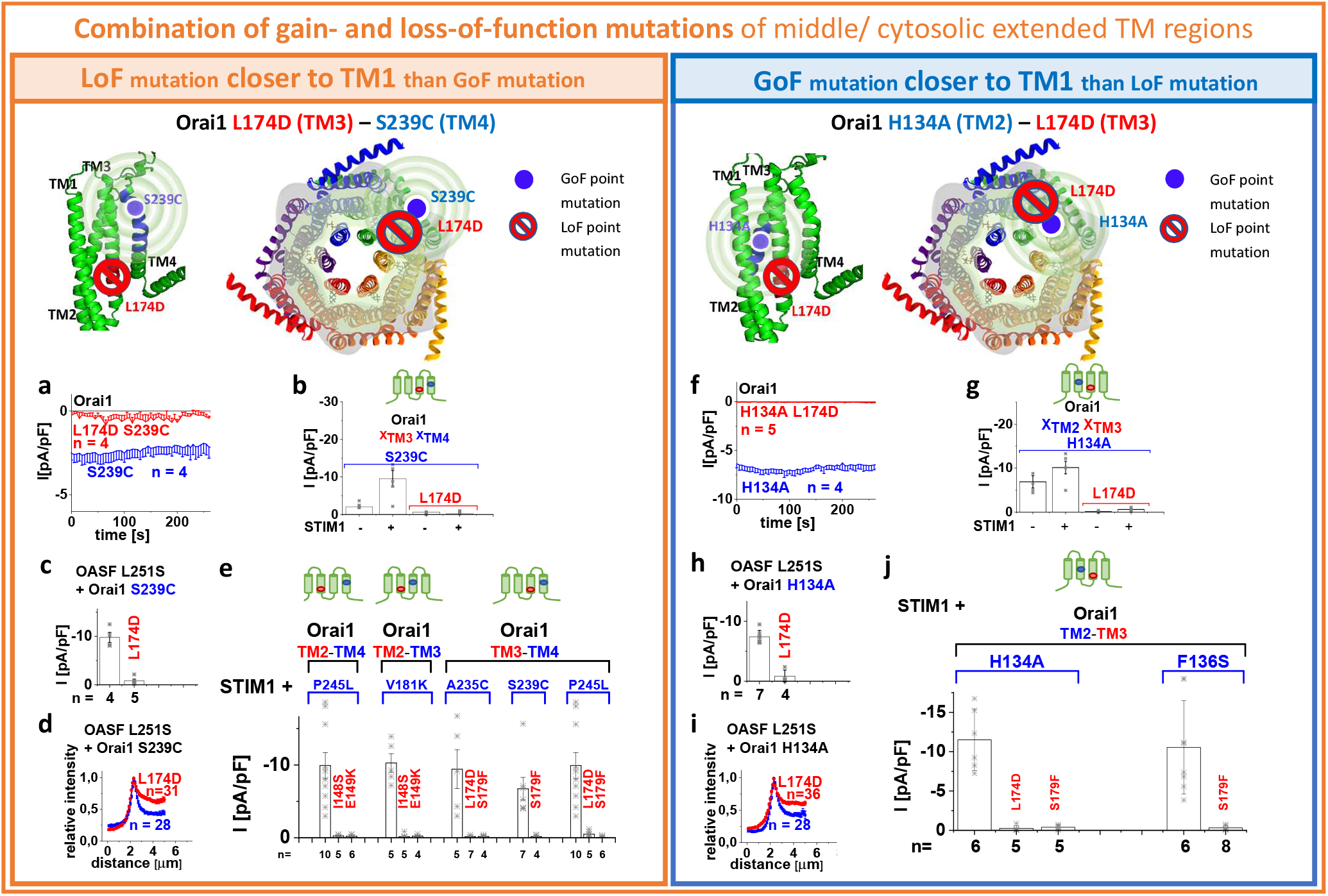
The combination of a GoF point mutation in the MTR with a LoF point mutation in the CETR leads to loss of function of the Orai channel independent of their position relative to each other. Four schemes representing the location of the investigated residues within a single subunit (top – left) of Orai1 or the whole channel complex (top – middle), for either LoF or GoF mutation closer to the pore, respectively. Red stop sign represents the position of the LoF mutation, while blue circle shows the position of the GoF mutation. The spheres indicate the impact of the GoF mutation on the entire subunit. In the following, it is valid for (a), (b), (c), (e), (f), (g), (h) and (j) that current densities of MTR-GoF-CETR-LoF double mutants are significantly different of those of the corresponding MTR-GoF mutants (p < 0,05). a) Time course of whole cell current densities of Orai1 S239C compared to Orai1 L174D S239C in the absence of STIM1. b) Block diagram of whole cell current densities of Orai1 S239C, Orai1 L174D S239C in the absence (t = 0 s) and the presence (maximum current densities) of STIM1 (n = 4 – 12 cells; values are mean ± SEM). c) Block diagram of whole cell current densities of Orai1 S239C compared to Orai1 L174D S239C in the presence of STIM1 OASF L251S. d) Intensity plots of STIM1-OASF-L251S co-expressed with Orai1 S239C compared to Orai1 L174D S239C (at 4 μm, p < 0,05). e) Block diagram of whole cell current densities of Orai1 P245L compared to Orai1 I148S P245L and Orai1 E149K P245L; Orai1 V181K compared to Orai1 I148S V181K and Orai1 E149K V181K; Orai1 A235C in comparison to Orai1 L174D A235C and Orai1 S179F A235C; Orai1 S239C in comparison to Orai1 S179F S239C; Orai1 P245L in comparison to Orai1 L174D P245L and Orai1 S179F P245L, all in the presence of STIM1. f) Time course of whole cell current densities of Orai1 H134A compared to Orai1 H134A L174D in the absence of STIM1. g) Block diagram of whole cell current densities of Orai1 H134A, Orai1 H134A L174D in the absence and the presence of STIM1 (n = 4 – 12 cells; values are mean ± SEM). h) Block diagram of whole cell current densities of Orai1 H134A compared to Orai1 H134A L174D in the presence of STIM1 OASF L251S. i) Intensity plots of STIM1-OASF-L251S co-expressed with Orai1 H134A compared to Orai1 H134A L174D (at 4 μm, p < 0,05). j) Block diagram of Orai1 mutant whole cell current densities of Orai1 H134A compared to Orai1 H134A L174D, Orai1 H134A S179F and Orai1 F136S compared to Orai1 F136S S179F, all co-expressed with STIM1.

In line with the findings above, we observed upon co-expression of STIM1 OASF L251S a total loss of function of both, Orai1 L174D S239C and Orai1 H134A L174D compared to the constitutively active, single point mutants Orai1 S239C and Orai1 H134A, respectively **(Fig 4 c, h)**. Investigation of the coupling of STIM1 via OASF L251S and OASF revealed significantly reduced, but not completely abolished, coupling of STIM1 to Orai1 L174D S239C and Orai1 H134A L174D **(Fig 4 d, i; Supp Fig 8 e, i).** Also, other double point mutants exhibit significantly reduced coupling to those STIM1 fragments as their corresponding constitutively active point mutants **(Supp Fig 8 b-d, g, h, l, o)**.

Also in this case, V102A in TM1 represents an exception. We discovered that none of the above reported, CETR-LoF mutations, as exemplarily tested with I148S, E149K and S179F were able to abolish the constitutive function of Orai1 V102A. These findings are also in line with published activity of Orai1 V102A L174D^49^. Co-expression of STIM1 was not able to enhance V_rev_ of V102A double mutants containing the cytosolic LoF point mutants **(Supp Fig 4 c-h)**, indicating that these CETR mutations hinder STIM1 coupling. In contrast, constitutive activity of two other GoF-TM1 mutants: F99M and V107M could be abolished upon the introduction of the CETR-LoF E149K mutation **(Supp Fig 4 i, j)**.

Summarizing, a series of MTR-GoF-CETR-LoF double point mutants showed that also LoF point mutations in the CETR act dominant over GoF point mutations, except for the GoF-V102A. Moreover, mutually dependent motions of the TM regions occur not only in one membrane plane within the MTR, but also across several membrane planes as here the MTR and CETR. Herewith, we provide fundamental evidence, that a conformational alteration of a single checkpoint residue spreads across the entire Orai1 subunit.

### LoF point mutations keep the pore of a constitutively active Orai1 mutant in a closed conformation

The above described dominant effect of diverse LoF over GoF mutations indicate that the LoF mutations keep the constitutively active Orai1 complex in the closed conformation. To verify that the Orai1 pore architecture of constitutively active mutants is affected by those LoF point mutations, we performed cysteine crosslinking and MD simulations on the following key double mutants Orai1 K85E H134A, Orai1 H134A E149K, Orai1 H134A L174D and Orai1 H134A S239W in comparison to Orai1 H134A. Frischauf et al.^46^ have recently demonstrated that cysteine crosslinking along the critical pore-lining residues, Orai1 R91C and Orai1 V102C, is significantly reduced upon the introduction of H134A. In contrast, crosslinking of those positions upon introduction of the double mutations: K85E H134A, H134A E149K, H134A L174D (only for Orai1 R91C) and H134A S239W (only for Orai1 R91C) exhibited again significantly enhanced levels of cysteine-crosslinking compared to the absence of the H134A mutation **(Fig 5 a, b)**.

**Figure 5:**
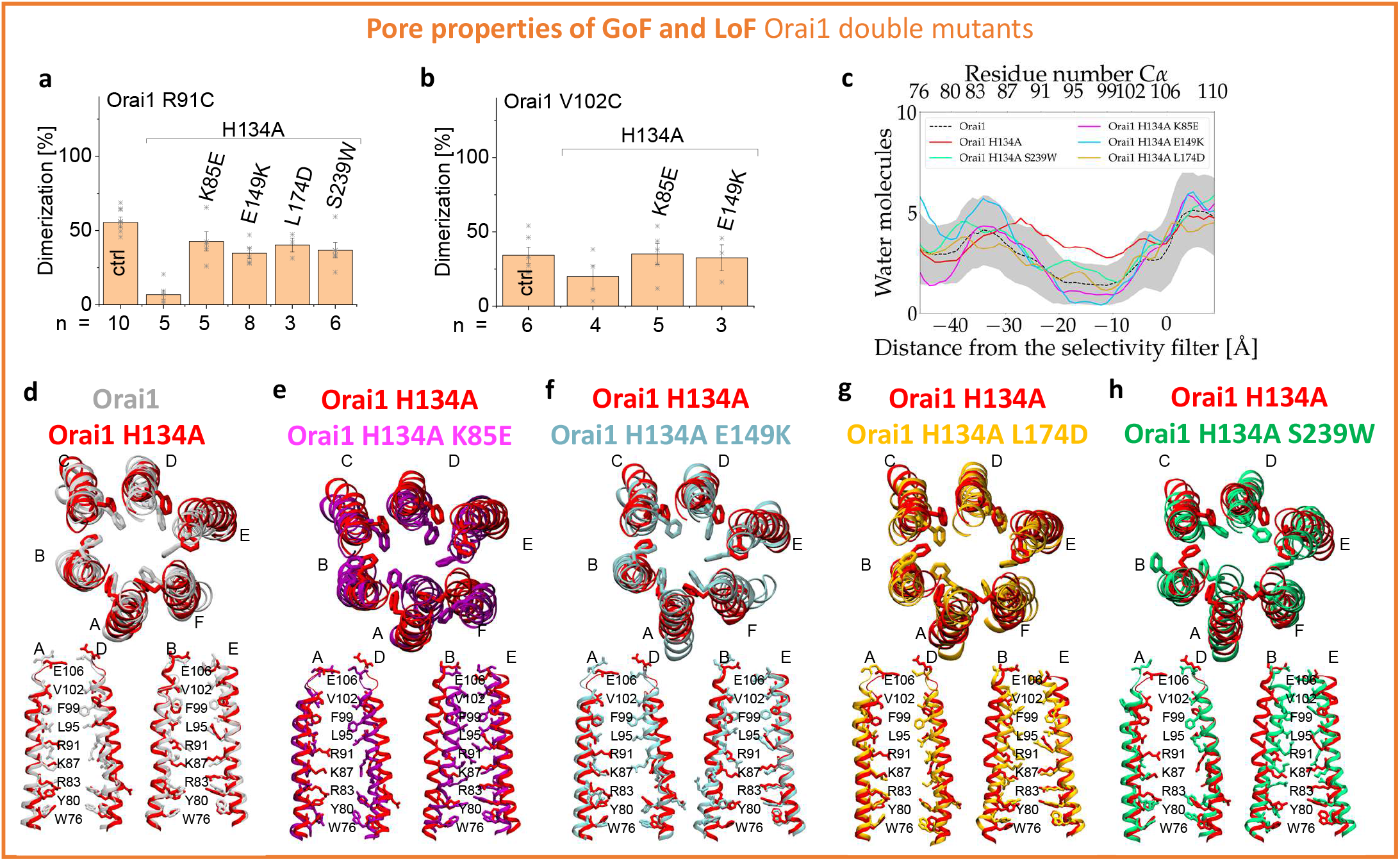
Orai1 MTR- and CETR-LoF point mutations hinder the formation of an opening-permissive pore geometry. a) b) Block diagrams exhibiting the degree of dimerization obtained via cysteine crosslinking of Orai1 R91C in comparison to Orai1 R91C H134A, Orai1 K85E R91C H134A, Orai1 R91C H134A E149K, Orai1 R91C H134A L174D and Orai1 R91C H134A S239W (a) and Orai1 V102C in comparison to Orai1 V102C H134A, Orai1 K85E V102C H134A and Orai1 V102A H134A E149K (b). Extent of crosslinking of single and triple mutants is significantly different compared to the double mutants. c) Pore hydration profile for wild type Orai1 and mutants. The number of water molecules is given as a function of the distance from the selectivity filter. Hydration profiles are given for wild type in dotted black line where the gray shaded areas correspond to the standard deviation of the mean. Solid red, purple, blue, orange and green lines correspond to Orai1 H134A, Orai1 K85E H134A, Orai1 H134A E149K, Orai1 H134A L174D and Orai1 H134A S239W. The profiles were calculated using the last 50 ns of the simulations. Positions of the carbon α of the residues delineating the pore are given on the top axis, while the distance from the selectivity filter is given on the bottom axis. (d) Molecular dynamics simulations demonstrate increased pore helix rotation and pore hydration in GoF mutation H134A. Superposition of snapshots at *t* = 350-400 ns from MD simulations of WT (gray) and H134A (red) as viewed from the top. Pairs of diagonal subunits viewed from the side (bottom). e – h) Superposition of snapshots at t = 350-400 ns from MD simulations of H134A (red) and K85E H134A (purple) (e), Orai1 H134A E149K (blue) (f), Orai1 H134A L174D (orange) (g), Orai1 H134A S239W (green) (h) as viewed from the top. Pairs of diagonal subunits viewed from the side (bottom).

In line with the findings by Yeung et al.^58^, our MD simulations up to 400 ns long revealed for Orai1 H134A in comparison to Orai1 wild-type enhanced hydration of the pore together with a rotation of F99 in the pore helix and a dilation of the pore. According to our functional data, the hydration of the pore of the double mutants Orai1 K85E H134A, Orai1 H134A E149K, Orai1 H134A L174D and Orai1 H134A S239W **(Fig 5 c)** was significantly reduced compared to Orai1 H134A. While Orai1 H134A L174D and Orai1 H134A S239W reached hydration levels comparable to Orai1 wild-type, Orai1 K85E H134A and Orai1 H134A E149K showed significantly enhanced dehydration also compared to Orai1 wild-type. Moreover, investigation of associated conformational changes revealed that F99 turned back to point again more into the pore region similar to wild-type Orai1 **(Fig 5 d-h; Supp Fig 9 a-f)**.

In summary, these data reveal that both CETR-as well as MTR-LoF mutations independent of their location relative to the GoF either in TM domains closer or farther apart and in distinct membrane planes (MTR, CETR) hamper pore dilation induced via a GoF mutation.

### Local enrichment of STIM1 overrules the dominant effect of MTR-, but not of CETR-LoF mutations

In the following, we tested whether local enrichment of STIM1 C-terminal fragments, induced by attaching two CAD fragments (-SS) to the Orai1 C-terminus of the double point mutants, is able to overcome the dominant role of LoF point mutations. Indeed, these close-by STIM1 fragments were able to overrule the dominant role of MTR-LoF over MTR-GoF point mutations, as visible by the constitutive activity of Orai1 H134W P245L – SS, Orai1 H134W V181K – SS and Orai1 H134A S239W – SS **(Fig 6 a-c; e)**.

**Figure 6:**
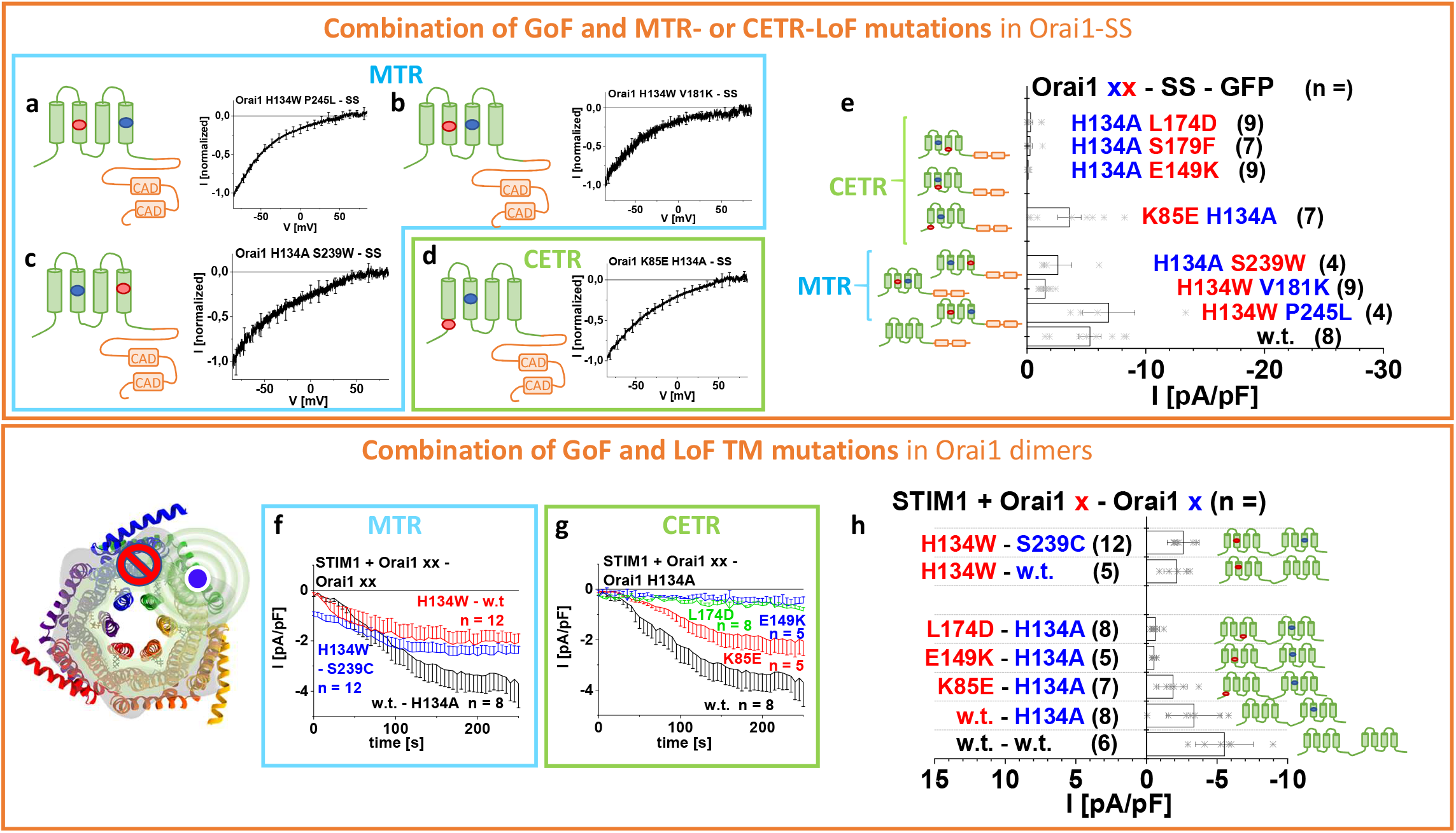
CETR-LoF mutations are dominant over MTR-LoF mutation both, upon local enrichment of STIM1-CAD and within an Orai1 dimer. a – d) Current/voltage relationships of Orai1 H134W P245L – SS-GFP, Orai1 H134W V181K – SS – GFP, Orai1 H134A S239W – SS – GFP and Orai1 K85E H134A – SS-GFP representing the effect of MTR-LoF mutations (blue; a-c) or the effect of CETR-LoF mutation (green; d). e) Block diagram of whole cell current densities of Orai1 – SS – GFP wild-type in comparison to Orai1 H134A L174D – SS – GFP, Orai1 H134A S179F – SS – GFP, Orai1 H134A E149K – SS – GFP, Orai1 K85E H134A – SS – GFP, Orai1 H134A S239W – SS – GFP, Orai1 H134W V181K – SS – GFP and Orai1 H134W P245L – SS – GFP. Current densities of mutants containing E149K, S179F or L174D are significantly different compared to the wild-type construct (p < 0,05). Green marked schemes represent the CETR LoF mutations, while blue labelled schemes show MTR-LoF mutations. Schemes display the location of one of the representative combinations of investigated residues within the whole Orai1 channel complex (bottom – left). Red stop sign represents the position of the LoF mutant, while blue circle shows the position GoF mutant. The spheres indicate the impact of the GoF mutation on the entire subunit and an adjacent subunit. f – g) Time course of Orai1 dimer whole cell current densities in the presence of STIM1. Orai1 dimer mutants represent Orai1 – Orai1 H134A compared to Orai1 H134W – Orai1 and Orai1 H134W – Orai1 S239C showing the effect of an MTR-LoF mutation (blue) (f). Orai1 dimer mutants represent Orai1 – Orai1 H134A in comparison to Orai1 K85E – Orai1 H134A, Orai1 E149K-Orai1 H134A and Orai1 L174D – Orai1 H134A showing the effect of a CETR LoF mutations (green) (g). h) Block diagram of whole cell current densities of Orai1 dimers in the presence of STIM1 corresponding to current densities recorded in f) and g) compared to the dimer formed by WT Orai1 (Orai1 – Orai1). Current densities of mutants containing E149K, S179F or L174D are significantly different compared to those of the wild-type construct (p < 0,05) a -e) & h) The schemes indicate the position of the corresponding mutations. Blue circle indicates GoF mutation, while red circle indicates the LoF mutation.

Also, the local attachment of two CAD fragments at the C-terminus of Orai1 K85E H134A (Orai1 K85E H134A – SS) is able to counteract the inhibitory effect of K85E even better as for Orai1-K85E-SS **(Supp Fig 6 i)**, which shows only small constitutive activity **(Fig 6 d, e)**. In contrast, double point mutants containing the MTR-GoF-H134A and another CETR-LoF mutation, remain inactive upon local attachment of STIM1 fragments at their C-termini (Orai1 H134A E149K – SS, Orai1 H134A S179F – SS, Orai1 H134A L174D – SS; **Fig 6 e**).

Overall, we discovered for MTR-GoF-LoF double mutants and Orai1 K85E-H134A that pore opening can still be re-established by enhancing local concentrations of STIM1. In contrast, other CETR-LoF point mutations seem to severely affect both, the pore geometry and STIM1 binding.

### CETR-LoF mutants are dominant over GoF mutants in Orai dimers

So far, we investigated the effects of LoF mutations in single Orai1 subunits. In the following, we were further interested whether above-described critical LoF mutations affect also GoF mutations of neighboring subunits.

Initially, we generated dimers with one wild-type subunit and one subunit, containing the constitutive MTR-GoF-H134A or -P245L mutation. In contrast to the strong constitutive activity of these point mutations in the homomer, in the heteromer, they allow only store-operated activation in the presence of STIM1 **(Fig 6 h; Supp Fig 10)** in line with the store-dependent activation of an Orai1 dimer containing one wild-type and one Orai1 V102A subunit^63^. This indicates that at least more than three GoF mutations within a hexamer are required to induce constitutive activity.

In the following, we generated a variety of Orai1 dimers containing an LoF mutation, in the first subunit and a GoF mutation in the second subunit. All of them, remained inactive in the absence of STIM1, while in the presence of STIM1 only some exhibited activation. In line with preserved STIM1 mediated activation of Orai1 H134W-Orai1, also Orai1 H134W-Orai1 S239C exhibited store-dependent activation upon co-expression with STIM1 to a similar extent as for Orai1-Orai1 H134A **(Fig 6 f, h)**. Interestingly, also Orai1 K85E-Orai1 H134A displayed store-operated currents **(Fig 6 g)**, in accordance with the activity of Orai1 K85E-Orai1 dimer published in Cai et al.^63^. In contrast, Orai1 E149K – Orai1 H134A and Orai1 L174D – Orai1 H134A remained inactive in the presence of STIM1 **(Fig 6 g, h)**.

To sum up, these data show that at least in the presence of STIM1 only CETR-LoF mutations except K85E act dominant over a GoF mutation in the neighboring subunit of a dimer and thus, overall in the whole channel complex in an allosteric manner. We demonstrate that inter-dependent motions initiated by an activation signal to induce pore opening spread not only in individual subunits, but also across neighboring subunits. The positions E149 and L174 are indispensable for this inter-dependent communication.

### Inter- and intra-Orai1 salt-bridge interactions in the CETR establish an intact STIM1 mediated Orai1 activation

Due to the more dominant role of the CETR-LoF mutations and to gain some insights into how K85E and E149K mutations can induce loss of function, we continued to investigate their role via MD simulations for selected mutants and extensive site-directed mutagenesis.

In MD simulations, we initially focused on the residues R83, K85, E149 and E173 and compared the axial positions of the carbon alpha relative to the hydrophobic gate as well as the distances between the charged moieties of each residue. The carbon alpha from K85 and E173 are on the same height, whereas the carbon alpha of R83 is located further toward the cytosol followed by the carbon alpha from E149 **(Supp Fig 11 a top)**. While the positions of the charged moieties from K85 and E173 follow the same observations, guanidine and carboxylate moieties from R83 and E149, respectively, are now at the same level as well **(Supp Fig 11 a bottom)**. However, the latter two residues are initially less likely to form salt-bridges as their side chains are oriented in the same directions **(Supp Fig 11 c)**. Despite this observation, even in its closed state, we noticed that TM1 offers enough flexibility in the basic region to allow for a slight rotation, allowing R83 to interact with E149 within one subunit **(Supp Fig 11 c).** Overall, we clearly observed that K85 and E173 within the same subunit and K85 and E149 of adjacent subunits have their side chains converging toward each other forming salt bridge interactions **(Fig 7 a, Supp Fig 11 b)**.

**Figure 7:**
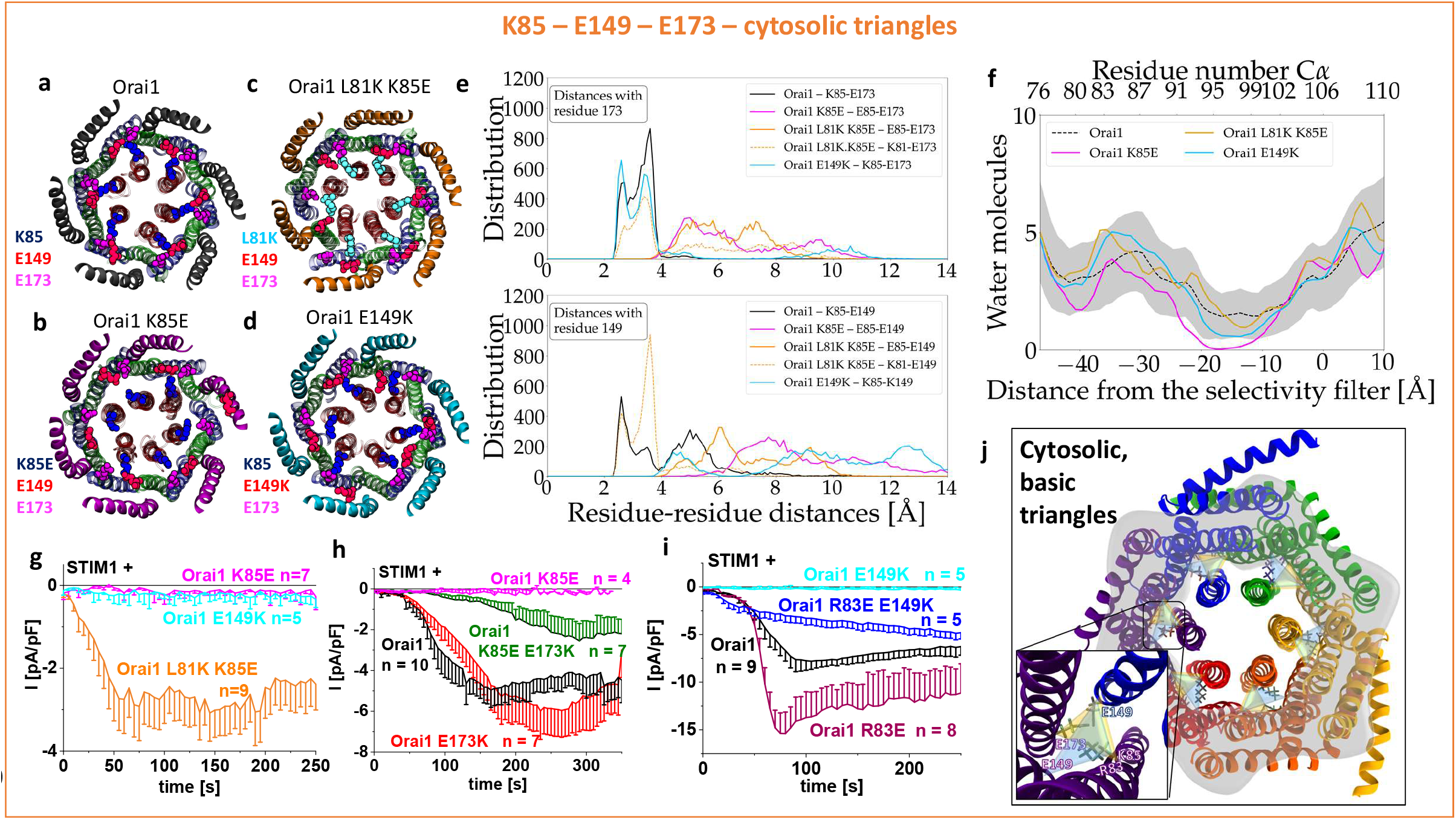
The cytosolic triangles contributes to the establishment of an opening-permissive pore geometry. a) – d) Cytosolic view of the starting and final snapshots of wild type Orai1 (a), Orai1 K85E (b), Orai1 L81K K85E (c) and Orai1 E149K (d) from molecular dynamics simulations. TM1, TM2 and TM3 are represented in glassy ribbon material in red, cyan and green respectively. TM4 is represented in a yellow opaque ribbon. Residues L81K, K85(E85), E149(K149) and E173 are colored cyan, blue, red and violet, respectively. Channel conformation is shown after 250 ns of molecular dynamics respectively. Differently colored C-termini mark the respective mutant with the color code also used in (e). e) Distribution of the distances calculated between the nitrogen atom of the lysine and the oxygen atoms from the carboxyl group of the glutamic acids for the last 50 ns of the simulations shown for the wild-type Orai1 (black), Orai1 K85E (purple), Orai1 L81K K85E (orange, dotted orange) and Orai1 E149K (cyan) (distance for E173 (top), distance for E149 (bottom)). Saltbridges were considered to be formed when the distance between the amino group of the lysine and the carboxyl group of the glutamic acids was below 5 Å. f) Pore hydration profile for wild type Orai1 and mutants. The number of water molecules is given as a function of the distance from the selectivity filter. Hydration profiles are given for wild type in dotted black line where the gray shaded areas correspond to the standard deviation of the mean. Dashed black, blue, purple and orange lines correspond to Orai1 wild-type, Orai1 E149K, Orai1 K85E and Orai1 L81K K85E. The profiles were calculated using the last 50 ns of the simulations. Positions of the carbon α of the residues delineating the pore are given on the top axis with distance from the selectivity filter shown on the bottom. g) Time course of STIM1-mediated whole cell current densities of Orai1 E149K, Orai1 K85E and Orai1 L81K K85E. h) Time course of STIM1-mediated whole cell current densities of Orai1 K85E, Orai1 E173K and Orai1 K85E E173K in comparison to wild-type Orai1. i) Time course of STIM1-mediated whole cell current densities of Orai1 R83E, Orai1 E149K and Orai1 R83E E149K in comparison to wild-type Orai1. j) The scheme with inset represents the two cytosolic triangles formed by the residues K85, E149, E173 and R83, E149, E173, respectively.

The K85E mutation led to the disappearance of these salt-bridge interactions in Orai1 **(Fig 7 b, e)**. K85E drifted slowly away from E149 and E173 **(Supp Fig 11 b)** which finally led to a collapse of the pore **(Fig 7 f)**. The hydration profiles **(Fig 7 f; Supp Fig 12 a-c)** corroborated these observations as the number of water molecules within the pore region of the channel significantly dropped in the case of Orai1 K85E. MD simulations on the Orai1 E149K revealed that especially the salt-bridge of K85-E149 was lost **(Fig 7 d, e)**, while the mean distance of K85-E173 slightly enlarged **(Supp Fig 11 a)**. This change was sufficient to also lead to a reduced amount of water molecules in the pore, concomitantly, with a clockwise rotation of F99 **(Supp Fig 12 a-c)**.

Introduction of a positively charged residue, L81K **(Fig 7 c, g)**, one helical turn N-terminal to K85E (Orai1 L81K K85E) restored, or at least, maintained the original pore structure since its hydration level is close to that of wild-type **(Fig 7 f)**. In line, the salt-bridge bonds with E149 of the neighbouring subunit and E173 within one subunit naturally occurring in wild-type and destroyed by the K85E mutation are restored in Orai1 L81K K85E **(Fig 7 c, e, f)**. Accordingly, a screen on several N-terminal mutations (L81K, S82K, A88K and S89K) revealed that STIM1 mediated activation and coupling of Orai1 K85E was reconstituted only by the additional introduction of L81K (Orai1 L81K K85E) **(Fig 7 g; Supp Fig 13 a).**

For further clarification of the impact of K85-E173 and K85-E149 salt-bridge interactions in Orai1 activation, we tested double mutants with the respective residues exchanged by ones with oppositely charged moieties. We uncovered that Orai1 K85E E173K recovers STIM1 mediated activation partially **(Fig 7 h)**, in line with Dong et al.^54^. In contrast, we find that Orai1 K85E E149K remains non-functional **(Supp Fig 13 b)**, despite our simulations clearly indicate a close distance and electrostatic interaction of K85-E149 **(Fig 7 a, e)**. The reason for that is probably that only in Orai1 K85E E149K, but not in Orai1 K85E E173K, the charged moieties at the respective positions move farther apart compared to wildtype. In search of further electrostatic interaction partners of E149, we identified R83 as a crucial residue. Orai1 R83E E149K displays restored STIM1 mediated activation **(Fig 7 i)**, in line with Dong et al.^54^ who suggested that R83 can rotate clockwise towards E149.

It is worth noting that among the functional relevant salt-bridge pairs R83-E149 and K85-E173, the two single point mutants Orai1 R83E and Orai1 E173K are functional in contrast to their respective counterparts K85E and E149K, despite in all single point mutants one salt-bridge is lost **(Supp Fig 13 b)**. Hence, we extended our functional screen to additional single and multiple point mutants containing diverse combinations of substitutions of these basic and acidic residues **(Supp Fig 13 b).** Further, we evaluated the number of potential attracting and repulsing forces of the residues at positions 83, 85, 149 and 173 **(Supp Fig 13 b)** in functional compared to non-functional mutants. This combined approach unveils that two intra-subunit salt-bridges: R83-E149, K85-E173 and one inter-subunit saltbridge K85-E149 play an essential role in maintaining Orai1 activation. Moreover, at least two of these communication pairs need to be intact to maintain Orai1 activation **(Fig 8 b, Supp Fig 13 b, Supp Fig 15 b)**.

**Figure 8:**
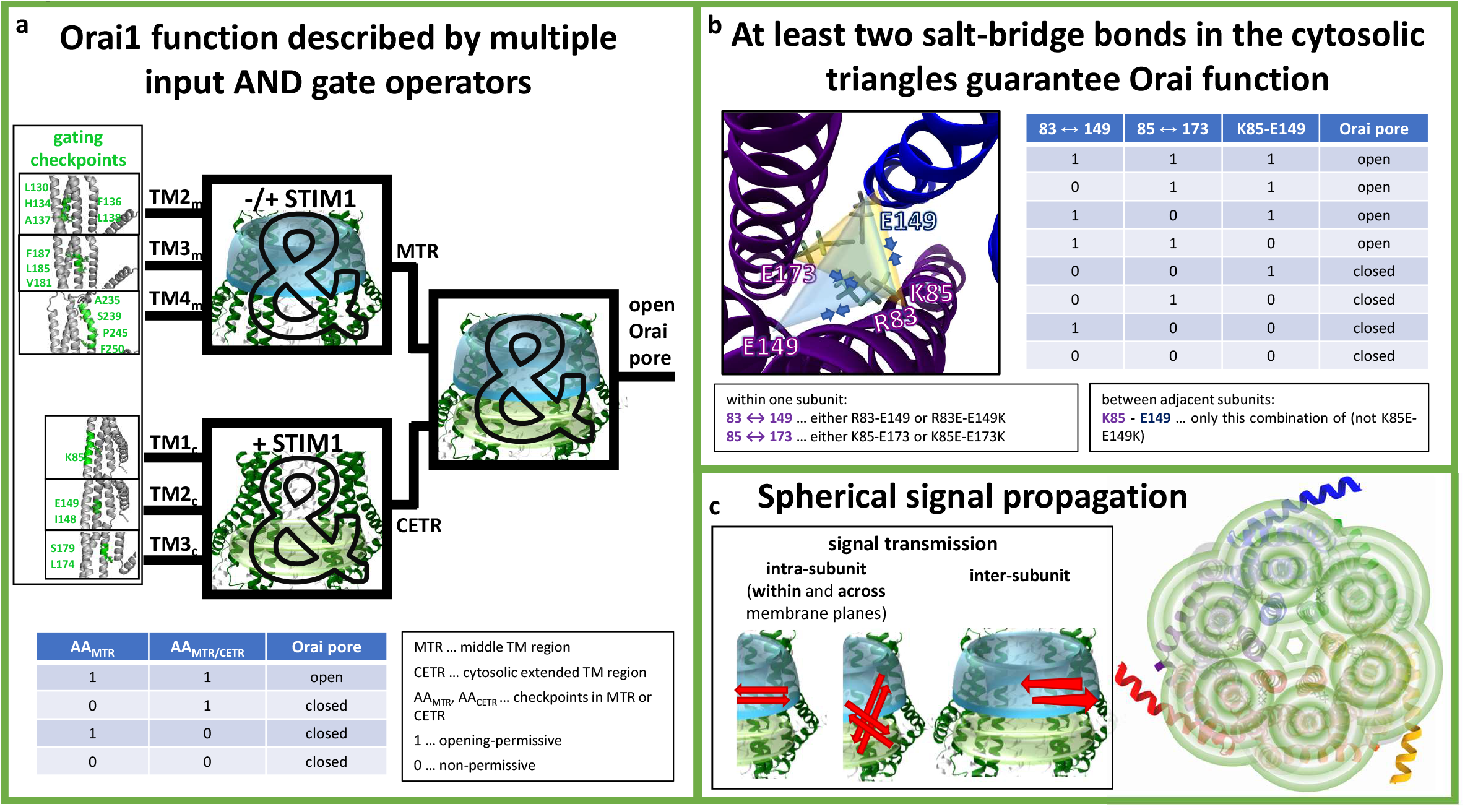
Orai1 signal propagation. a) Orai pore opening requires a series of gating checkpoints to capture an opening-permissive conformation. A single residue in a non-permissive conformation impairs pore opening. These complex correlations can be best described by the operator symbol for the AND gate containing multiple inputs, used in digital electronics. These multiple inputs include gating checkpoints in the MTR and the CETR. Each section individually, but also both together, can be viewed as an AND gate. The truth table (bottom), simplified for just two gating checkpoints AA_MTR_ and a second AA_MTR_ or an AA_CETR_ shows that as soon as one of two gating hots adopts a non-permissive conformation, the Orai channels remain closed. b) The CETR includes the cytosolic triangles which form salt-bridge interactions within and between subunit(s). Three pairs of charged residues R83-E149, K85-E173 and K85-E149 are functionally relevant. Among those at least two salt-bridges are required to be intact and maintain STIM1-mediated Orai1 activation. These requirements can be described by the truth table in (b) and thus their interplay can be illustrated by an OR gate of always two salt-bridges (see Supp Fig 14 b) that maintain an opening-permissive channel conformation. To maintain Orai1 function only the pairs R83-E149 and K85-E149, but not the K85-E149, can be mutated to oppositely charged residues. c) An Orai1 activation signal either from a constitutively active mutation or from STIM1 coupling to Orai1 C-terminus impacts each subunit (via alteration of intra-subunit interactions) within and across membrane planes (MTR, CETR)) and thus, the entire channel complex (via alteration in inter-subunit interaction).

In accordance with the functional data, we discovered that the salt-bridge interactions in the CETR also impact STIM1 coupling. While the plasma membrane intensity of the STIM1 C-terminal fragments, OASF and OASF L251S, was strongly reduced when co-expressed with these single point mutants K85E and E149K, the double point mutants Orai1 K85E E173K and Orai1 R83E E149K displayed again restored coupling of STIM1 **(Supp Fig 12 e-g).**

In conclusion, the sophisticated interplay of the basic and acidic residues in the CETR is a core element for CRAC channel function, pore hydration, pore opening and STIM1 coupling. We termed these crucial salt bridge interactions established within and between adjacent subunits, the cytosolic triangles.

### Orai1 CETR-LoF L174D at the hinge plate does not interfere with the cytosolic triangle

Orai1 L174D, a recently reported LoF mutant in the hinge plate^49^, has been proposed to affect the communication with L261 in TM4 and in consequence signal propagation to the pore. As shown in **Supp Fig 14 a,** MD simulations revealed that L174D affects the hydrophobic gate of Orai1 in a similar way to K85E and E149K.

In a further step, we investigated whether in Orai1 L174D, the negatively charged aspartate, interferes with the salt bridge interactions between TM1-TM2-TM3, due to its close positioning to E173. However, MD simulations revealed that Orai1 L174D widely maintains the K85-E173 and K85-E149 salt bridge interactions both, in Orai1 and Orai1 H134A **(Supp Fig 14 b)**. Our following data indicate in line with Zhou et al.^49^ that L174 in TM3 maintains Orai1 function via communication with L261 in TM4. Though we were unable to restore STIM1 mediated Orai1 activation via the introduction of oppositely charged residues at L174 and L261 (Orai1 L174D L261K, Orai1 L174K L261D; **Supp Fig 14 c right**), we discovered that a double point mutant Orai1 E173K L261D displays pronounced activity in contrast to strongly reduced currents of Orai1 L261D **(Supp Fig 14 c)**. A triple mutant Orai1 K85E E173K L261D displays again a loss of store-operated activation in the presence of STIM1 **(Supp Fig 14 c right)**, despite maintained function of Orai1 K85E E173K **(Fig 7 h)**.

In summary, besides the triangular interaction of basic and acidic residues in the CETR, the interplay of hydrophobic leucines in TM3 and TM4 represent a second individual module required for an intact Orai1 channel. The L174D mutation affects the pore in a similar manner like Orai1 K85E and Orai1 E149K, but does not interfere with the salt-bridge interactions of the cytosolic triangles.

## Discussion

In this study we provide profound evidence for the previously proposed wave of inter-dependent motions of TM helices^51^ within the Orai channel upon pore opening. This assumption of a global concerted gating motion is based on the recent discovery of a series of GoF mutants throughout all TM domains^51^. Our library of GoF-LoF double point mutants **(Table 1)** unveils a dominant inhibitory effect of a series of LoF over most GoF mutations independent of their distance to the pore and their location relative to each other **(Table 1)**. This suggests that global instead of local conformational changes within the entire Orai complex are fundamental for pore opening. We discovered that these gating motions are controlled by crucial checkpoints, located predominantly in the conical MTR-ring, the CETR including the cytosolic triangles. LoF mutations at these key positions interfere mainly with an open pore geometry. As long as clearance of all these critical checkpoints is achieved pore opening of the CRAC channel complex occurs. On the contrary, a single position in a non-permissive state inhibits Orai channel opening. Thus, these complex correlations can be best described by the principles of Boolean algebra. We show that the Orai pore opening mechanism in dependence of gating checkpoints in the MTR and CETR follows the rules of the truth table of a logical “AND” gate as shown in **Fig 8 a; Supp Fig 15**.

A significant feature of several residues in the conical MTR ring is that their single point mutation can lead either to gain-(MTR-GoF) or loss-of-function (MTR-LoF). Thus, these residues possess two roles: (1) maintaining the closed state of the Orai channel, as mutations can lead to constitutive activation likely due to the release of steric brakes as recently studied in detail for the position H134^46,58^ and (2) contributing to the establishment of an opening-permissive pore and channel conformation, as other mutations can abolish function likely due to a dewetting of the hydrophobic gate.

Critical checkpoints located within the CETR, in particular in the cytosolic triangles and the hydrophobic modules, establish an opening-permissive pore geometry as revealed by cysteine crosslinking and MD simulation studies. Within the cytosolic triangles of two adjacent subunits, two intra-Orai1 and one inter-Orai1 salt-bridge interactions formed by basic and acidic residues in TM1, TM2 and TM3 are relevant in establishing an opening-permissive pore conformation. While the available closed and open Orai structures indicate that two of them (K85-E173, K85-E149) are already in close proximity, one (R83-E149) seems to require larger rearrangements to form a stable salt-bridge. K85 in Orai1 appears to be a critical pivot for the channel through its interactions with E149 and E173. These triangular interactions possess probably two major functions: (1) They prevent the pore from collapsing and (2) they keep E149 and E173 close to each other in preparation for further conformational reorientations during pore opening after STIM1 binding. The latter refers for instance, to the functional importance of the R83-E149 salt-bridge **(Fig 8 b).** We suppose that it is preferentially formed in the presence of STIM1. In line with this hypothesis, Dong et al.^54^ suggested that upon channel opening rotation in TM1 might happen. Then, R83 can get in contact with E149 which is kept in the vicinity due to its interaction with K85 allowing the pore to further expand. This new interaction seems to act as a second lock, similar to K85-E173, to keep the channel in a fully open state. Moreover, we assume that it assists to stretch the hydrophobic gate even further to increase the level of hydration, which facilitates the permeation of Ca^2+^ ions through the channel. Our functional studies further elucidated that at least two of these three salt-bridges are required to be intact for a functional pore conformation and the maintenance of STIM1-mediated Orai1 activation **(Fig 8 b)**. Interestingly, these correlations can be illustrated by the logical operators for the AND and OR gates as shown in **Supp Fig 15.**

Our pool of double point mutants containing one MTR-GoF and one MTR- or CETR-LoF in a variety of combinations provide one key step further in the detailed understanding of the motions within the Orai complex upon pore opening. A diversity of LoF mutations prevent a GoF mutant to switch into the open state, likely via the introduction of inter-TM constraints. Thus, we propose that the Orai1 activation signal, induced by STIM1 coupling to the Orai1 C-terminus, propagates like a spherical wave of inter-dependent TM motions within one subunit to the entire hexameric channel complex **(Fig 8c)** to induce pore opening. This concept is especially supported by three observations: (1) Double mutants containing a GoF and a LoF point mutation, independent of which one located in the TM domain closer to the pore, lose function. (2) LoF mutations are not only dominant over GoF mutations within the same (e.g. MTR-LoF and MTR-GoF), but also over distinct membrane planes (e.g. CETR-LoF over MTR-GoF such as L174D over H134A; **Fig 8c; Table 1)**. (3) The dominant inhibitory effect holds also for some combinations of LoF (E149K, L174D) and GoF mutations, when each is located within a distinct subunit of a dimer. Concerning the latter, the requirement of more than three intact Orai subunits is in line with the non-linear dependence of Orai activation and STIM1 coupling^64^. The transmission of the activation signal across the entire Orai channel complex is likely enforced by the TM2/3 ring which has recently been proposed to function as an individual unit that devolves strong cooperativity between STIM1 coupling at six single subunits and pore opening^58,65,66^.

Among constitutively activating Orai1 point mutations, the V102A in TM1 is the only GoF mutation that acts dominantly over MTR- and CETR-LoF point mutations. These findings suggest that the extent of pore hydration of Orai1 V102A^53^ is sufficient to confer Ca^2+^ permeation. This exceptional behavior of Orai1 V102A TM1 pore mutant is also accompanied by a number of distinctly different biophysical hallmarks distinguishing it from the TM2-TM4-constitutive point mutants. Only Orai1 V102A leads to a strong reduction of V_rev_, an enhancement of I_DVF_ versus I_Ca2+_, allows the permeation of Cs^+^ ions and maintains the activity upon N-terminal deletions^32,36,46^. Thus, Orai1 V102A activation occurs via pronounced pore dilation or distinct inter-dependent motions of the TM helices, which is likely responsible for its unique biophysical characteristics^36^ that are so different to other Orai activating mutants.

Whether the global conformational changes within the Orai complex upon pore opening are provoked via the STIM1 coupling to Orai1 C-terminus or additional other cytosolic domains is still unclear. At present, one hypothesis is that STIM1 coupling to the Orai1 C-terminus is sufficient to trigger the allosteric movements within the channel complexes^49^. An alternative hypothesis is that the interdependent TM movements are additionally triggered by STIM1 coupling to the loop2 and the N-terminal segments^34^. The effect of LoF mutations enabled us to elucidate sites critical for STIM1 coupling and to clarify their significance relative to each other. Only the inhibitory effect of MTR-LoF and the CETR-K85E mutations can be partially overcome by co-expression of OASF L251S with Orai1 or by the local attachment of CAD fragments direct onto the respective Orai protein. Moreover, dimers containing one subunit with a GoF mutation and the second subunit with either an MTR-LoF or the CETR-K85E mutation remain functional. This indicates that an opening-permissive pore geometry is retained or at least can be forced by an adequate amount of STIM1 bound to Orai channels. In contrast, other CETR-LoF mutations (e.g. E149, L174) prevent these alternative activation pathways. In line with these findings, co-localization studies revealed that the MTR-LoF and the CETR-LoF Orai1 K85E mutations affect STIM1 coupling only mildly, while other CETR-LoF mutations impede STIM1 coupling. This behavior is also reflected by the V_rev_ of the respective LoF mutants in the V102A background, thus functioning as a readout parameter for STIM1 coupling^30,32^. Hence, gating checkpoints in or close to the loop2 region seem to contribute to the formation of a STIM1 coupling site, either directly or indirectly. Accordingly, the loop2 segment has been suggested to be involved in STIM1-mediated gating^35^. Moreover, the N-terminus together with the loop2 region is required to establish an opening-permissive pore conformation and to fine-tune authentic CRAC channel hallmarks together with STIM1^34^. Thus, the loop2 with E149 and L174 is besides Orai1 C-terminus, the second indispensable prerequisite for a fully intact STIM1-Orai1 coupling. Due to the more potent inhibitory role of E149K and L174D compared to K85E, it is tempting to speculate that the loop2 region functions as an essential bridge for transmission of the Orai1 activation signal from the Orai1 C-terminus to the pore.

In addition to the indispensable checkpoints in the MTR and CETR, the MTR harbors several positions only crucial for STIM1 mediated activation, but not for H134A induced activation and STIM1 binding as determined in Fig 4 a and Supp Fig 4 a-d. This suggests partly distinct requirements in the signal propagation pathways of STIM1 and H134A mediated activation. Moreover, the inhibitory action to STIM1 coupling and -mediated activation of LoF mutations in the hinge region (3xA,3xG, L261D) can be bypassed by MTR-GoF mutations (H134A, P245L). Thus, we assign to the hinge region a potent role in STIM1 coupling in line with Zhou et al.^49^, besides its potential effect on the pore architecture.

Overall, we prove that Orai1 pore opening comes along with opening-permissive, inter-dependent TM domain motions across the entire channel complex. This requires clearance of a number of gating checkpoints in the middle and cytosolic segments of the Orai TM domains. A core element for STIM1 coupling and Orai1 pore opening is established by cytosolic triangular salt-bridge interactions within one and between neighboring Orai1 subunits. A synergistic action of these gating checkpoints together with STIM1 coupling triggers pore opening and ensures high Ca^2+^ selectivity. The variety of elucidated mutants represent promising tools in future drug development to interfere with one of the multiple gating checkpoints to manipulate Orai1 function. Moreover, they provide a valuable source for novel structural insights.

## Experimental Procedures

### Molecular Biology

For N-terminal fluorescence labeling of human Orai1 (Orai1; Accession number NM_032790_3, provided by A. Rao’s lab) the constructs were cloned into the pEYFP-C1 (Clontech) expression vector via KpnI and XbaI (Orai1) restriction sites. Site-directed mutagenesis of all the mutants was performed using the QuikChange™ XL site-directed mutagenesis kit (Stratagene) with the corresponding Orai1 constructs serving as a template.

Human STIM1 (STIM1; Accession number: NM_003156) N-terminally ECFP-tagged was kindly provided by T. Meyer’s Lab, Stanford University. pECFP-C1 STIM1 C terminus (aa233-685 wt and L251S) was used as a template for the generation of pECFP-OASF (wt and L251S) by introducing a stop codon at position 475 (aa 233–474) using the QuikChange XL site-directed mutagenesis kit (Stratagene). STIM1 fragment 344-449 (CAD) was amplified via PCR including an N-terminal KpnI and a C-terminal XbaI restriction site for cloning into the pECFP-C1 vector.

### Orai1 dimers with a linker

Point mutations (H134A / S239W / K85E / E149K / L174D) were introduced into peYFP-C1/Orai1 (peYFP-C1 – Clontech) plasmids. YFP-Orai1 dimers were cloned using suitable restriction sites. Linker regions (RDPLVQGGGSGGCGGIAL) between Orai1 cDNAs were constructed using QuikChange site-directed mutagenesis kit (Agilent Technologies) by the use of suitable primers inserting amino acids GGGSGG in dimers that included the short linker region (RDPLVQ*CGGIAL).

### Orai1-CAD-CAD-constructs

Orai1-CAD-CAD-GFP constructs were designed by cloning Orai1 and STIM1-CAD domains (aa344-449) into peYFP-N1 (Clontech). Linker regions between Orai1 and CAD (Orai1-linker-1-CAD-linker-2-CAD-YFP) domains were constructed using QuikChange site-directed mutagenesis with suitable primers (linker-1: SSRAGGGGSGGGGS; linker-2 GGSGGGSSPRGGGGGSGGGGS) inserting desired amino acids.

The integrity of all resulting clones was confirmed by sequence analysis (Eurofins Genomics/Microsynth).

### Cell culture and transfection

The transient transfection of HEK293 cells was performed^67^ using the TransFectin Lipid Reagent (Bio-Rad) (New England Biolabs).

The transient transfection of HEK293 cells was performed^67^ using the TransFectin Lipid Reagent (Bio-Rad) (New England Biolabs). Regularly, potential cell contamination with mycoplasma species was tested using VenorGem Advanced Mycoplasma Detection kit (VenorGEM).

### Cell preparation, cysteine-crosslinking and western blot analysis

HEK293T cells cultured in 12 cm dishes were transfected with 5 μg plasmid using Transfectin lipid reagent (Biorad) following the manufacturer’s instructions. 24 hours after transfection, cells were harvested and washed twice in an HBSS (Hank’s balanced salt solution) buffer containing 1 mM EDTA.

After centrifugation (1000 g/2 min), cell pellets were resuspended in homogenization buffer [25 mM Tris HCl pH 7.4, 50 mM NaCl, protease inhibitor (Roche)] and incubated on ice for 15 min. Lysed cells were passed 10 times through a 27G ½” needle and centrifuged at 1000 g for 15 min at 4°C to pellet debris. Cell lysates were treated with 1mM CuP-solution for 5 minutes at room temperature and the reaction was stopped upon addition of 50 mM N-ethylmaleimid quenching solution. 21μl of each sample was mixed with nonreducing Laemmli’s buffer, heated 15 min at 55°C, and subjected to a 12% SDS PAGE. Separated proteins were transferred to a nitrocellulose membrane and immunoblotted with an antibody recognizing Orai1 (Sigma Aldrich). Each experiment was performed at least 3 independent times and analysed with the program ImageJ (NIH) to calculate the percentage of dimer formation.

### Electrophysiology

Electrophysiological recordings that assessed the characteristics of 2-3 constructs were carried out in paired comparison on the same day. Expression patterns and levels of the various constructs were carefully monitored by confocal fluorescence microscopy and were not significantly changed by the introduced mutations. Electrophysiological experiments were performed at 20-24°C, using the patchclamp technique in the whole-cell recording configuration. For STIM1/Orai as well as STIM1 C-terminus/Orai current measurements, voltage ramps were usually applied every 5 s from a holding potential of 0 mV, covering a range of −90 to +90 mV over 1s. The internal pipette solution for passive store-depletion contained (in mM) 3.5 MgCl_2_, 145 Caesium Methane Sulphonate, 8 NaCl, 10 HEPES, 20 EGTA, pH 7.2. Extracellular solution consisted of (in mM) 145 NaCl, 5 CsCl, 1 MgCl_2_, 10 HEPES, 10 glucose, 10 CaCl_2_, pH 7.4. Applied voltages were not corrected for the liquid junction potential, which was determined as +12mV. All currents were leak-corrected by subtraction of the leak current which remained following 10 μM La^3+^ application.

Bar graphs in the figures display for Orai1 proteins in the absence of STIM1 the current density at t = 0 s, while in the presence of STIM1 maximum current densities are shown.

### Confocal fluorescence microscopy

Confocal microscopy for co-localization experiments was performed similarly to^68^. In brief, a QLC100 Real-Time Confocal System (VisiTech Int., UK) was used for recording fluorescence images connected to two Photometrics CoolSNAPHQ monochrome cameras (Roper Scientific, USA) and a dual port adapter (dichroic: 505lp; cyan emission filter: 485/30; yellow emission filter: 535/50; Chroma TechnologyCorp., USA). This system was attached to an Axiovert 200M microscope (Zeiss, Germany) in conjunction with an argon ion multi-wavelength (457, 488, 514 nm) laser (Spectra Physics, USA). The wavelengths were selected by an Acousto Optical Tuneable Filter (VisiTech Int., UK). MetaMorph 5.0 software (Universal Imaging Corp.) was used to acquire images and to control the confocal system. Illumination times for CFP and YFP images that were consecutively recorded with a minimum delay were about 900 msec.

### Simulation protocols

A homology model of hOrai1 based on the crystal structure (34) was used through all the simulations following the procedure described in Frischauf *et. al.* (62). Both wild type and Orai mutants were embedded in a pure POPC membrane composed by 388 lipids using the CHARMM-GUI web interface (63). The TIP3p (64) model is used to represent water molecules. The CHARMM36 forcefield was used to treat protein (65) and lipids (66). Calcium and chloride ions were used to neutralize the system with an ionic strength set at 100 mM. One calcium ion was placed in the selectivity filter accordingly to the crystal structure prior running any simulations. Ions were treated by using a rescaled charge paradigm (67-70). These parameters have been shown to better reproduce the properties of the ion in water and its hydrated structure (70). Interactions with the backbones of proteins and the free-energy of binding of Ca^2+^ ions to Ca^2+^-signaling proteins (71) are also improved.

Mutations were performed using the following protocol. A wild type Orai configuration was extracted from a 200 ns long simulation. The *in silico* mutation tool of the modeling program Yasara (72) was used to introduce the mutations.

GROMACS (73) software version 5.1.4 was used to perform all MD simulations. The temperature was set at 310K under a pressure of 1 atm. Minimization and equilibration followed the six-steps CHARMM-GUI protocols (74,75). 250ns and 400ns long molecular dynamics production simulations were performed for single and double point mutants, respectively.

VMD (76) was used to visualize the simulations. VMD and MD analyses (77) were used to analyze the simulations. Figures were generated using Matplotlib (78). The last 50 ns of each trajectory were used for analysis.

### Statistics

Results are presented as means ± s.e.m. calculated for the indicated number of experiments. The Student’s two-tailed t-test was used for statistical comparison considering differences statistically significant at p < 0.05.

## Acknowledgements

We thank S. Buchegger for excellent technical assistance. This work was supported in part by the Linz Institute of Technology project LIT-2018-05-SEE-111 and the Austrian Science Fund (FWF) projects P27641, P30567, P32851 and to I.D., P32947 to M.F., P32075-B to I.F, P28701 to R.S., P27263, P33283 and PhD program W1250 “NanoCell” to C.R. D.B. was supported the Czech Science Foundation (19-20728Y). Access to the National Grid Infrastructure Metacentrum and provided computational resources are gratefully acknowledged.

## Author contributions

AT and ID conceived and coordinated the study and wrote the paper. ID, AT, CH, MS performed and analyzed electrophysiological experiments. MS and RS carried out fluorescence microscopy experiments. CH, VL, MS, IF, MF, SL, LM, SB contributed to molecular biology and biochemistry. VL and IF performed cysteine crosslinking studies. DB, SP and RE contributed to the MD simulations. All authors reviewed the results and approved the final version of the manuscript.

