## Supplementary Figures for "A series of Orai1 gating checkpoints in transmembrane and cytosolic regions requires clearance for CRAC channel opening: Clearance and synergy of Orai1 gating checkpoints controls pore opening"

**Supplementary Figure Legends:**

**Supplementary Figure 1: Screen across residues in TM2, TM3 and TM4 in close proximity.** a) – f) Block diagrams depicting maximum current densities of Orai1 mutants in TM2, TM3 and TM4 in the absence (a – c) and presence (d – f) of STIM1. Only those positions which show close proximity (2-4 Å) to a residue in an adjacent TM domain as depicted in Table 2 and 3. Constitutive mutants (defined as > 0,5 pA/pF) are marked with a blue ball. Loss-of-function mutants (defined as < 0,5 pA/pF) are marked with a red stop sign.

**Supplementary Figure 2I and 2II: Activity of TM2, TM3 and TM4 mutants in dependence of degree of hydrophobicity (I) and of the size of the introduced amino acid (II).** a) – r) Current densities of L130X (a, b), F136X (c, d), V181X (e, f), L185X (g, h), F187X (i, j), A235X (k, l), S239X (m, n), P245X (o, p) and F250X (q, r) mutants in the absence compared to the presence of STIM1 plotted against the hydrophobicity (Supp Fig 1I) or the size of the introduced amino acid (Supp Fig 1II) .

**Supplementary Figure 3: Properties of Orai1 GoF and LoF mutants in the MTR.** a) Intensity of plasma membrane fluorescence of YFP-labelled Orai1 H134A, S239C, S239W, H134W S239C and Orai1 H134A S239W. b) - d) Intensity plots of STIM1-OASF co-expressed with Orai1 V181K compared to Orai1 H134W V181K (b), Orai1 P245L compared to Orai1 H134W P245L (c) and Orai1 S239C compared to Orai1 H134W S239C (d). e) Intensity of plasma membrane fluorescence of YFP-labelled Orai1 V181K, Orai1 H134W V181K, Orai1 P245L and Orai1 H134W P245L. f) g) Intensity plots of STIM1-OASF co-expressed with Orai1 H134A compared to Orai1 H134A A235W (f) and Orai1 H134A compared to Orai1 H134A S239W (g). h) Intensity of plasma membrane fluorescence of YFP-labelled Orai1 H134A, Orai1 A235W and Orai1 H134A A235W. i) Block diagram of current densities of Orai1 P245L compared to Orai1 P245L A235W. j) Intensity plots of STIM1-OASF co-expressed with Orai1 P245L compared to Orai1 A235W P245L. k) Intensity of plasma membrane fluorescence of YFP-labelled Orai1 P245L, Orai1 A235W and Orai1 A235W P245L.

**Supplementary Figure 4: Orai1 mutant currents in the background of V102A or a hinge mutation.** a) c) e) g) Current/voltage relationship of Orai1 V102A H134W, Orai1 V102A I148S, Orai1 V102A E149K and Orai1 V102A S179F in the absence compared to the presence of STIM1. b) d) f) h) $V_{rev}$ ($V_{rev}$ represents the reversal potential) of the respective mutants in a) c) e) g). i) j) Time course and corresponding block diagram of STIM1 mediated current densities of Orai1 F99M compared to Orai1 F99M H134W, Orai1 F99M E149K and Orai1 V107M compared to Orai1 V107M H134W and Orai1 V107M E149K. k) Block diagram of whole cell current densities of the Orai1 hinge mutants: Orai1 3xA and Orai1 3xG in the presence of STIM1 compared to Orai1 3xA H134A, Orai1 3xA P245L, Orai1 3xG H134A and Orai1 3xG P245L in the absence and the presence of STIM1. l) Intensity plots of STIM1-OASF co-expressed with Orai1 3xG compared to Orai1 3xG H134A and Orai1 3xG P245L. m) Block diagram of whole cell current densities of the Orai1 L261D in the presence of STIM1 compared to Orai1 P245L L261D in the absence and presence of STIM1. n) Block diagram of whole cell currents of the Orai1 ANSGA compared to Orai1 H134W ANSGA and Orai1 A235W ANSGA in the presence of STIM1.

**Supplementary Figure 5: Novel loss of function point mutations in the MTR.** Scheme representing the location of the MTR region (cyan) and the CETR (green) in the whole channel complex. a) Time course of STIM1 mediated MTR LoF Orai1 L188S, Orai1 L194S and Orai1 M243S current densities. STIM1 mediated current densities of the LoF mutants are significantly different compared to those of Orai1 wild-type. b) Intensity plots of STIM1-OASF co-expressed with Orai1 mutants shown in (a) compared to wild-type Orai1 (at 4 µM, $p < 0,05$ for MTR mutants). c) Block diagram of Orai1 double mutant current densities including Orai1 H134A T142C, Orai1 H134A L188S, Orai1 H134A V191N, Orai1 H134A L194S and Orai1 H134A M243S in comparison to Orai1 H134A in the absence of STIM1. d) Block diagram of STIM1 mediated Orai1 double mutant current densities including the mutants from c) Orai1 in comparison to Orai1 H134A ($p < 0,05$).

**Supplementary Figure 6: Activity of Orai1 mutants in the CETR.** a) Intensity of plasma membrane fluorescence of YFP-labelled Orai1 I148S, Orai1 E149K, Orai1 S179F, Orai1 L188S and Orai1 L194S. b) Block diagram of STIM1 mediated whole cell currents of Orai1 wild-type compared to Orai1 T142C and Orai1 T142C. c) Current densities of the S179X mutant in the presence of STIM1 plotted against the hydrophobicity. d) Block diagram of STIM1 mediated whole cell currents of Orai1 wild-type compared to Orai1 E149K, Orai1 E149D and Orai1 E149A.

**Supplementary Figure 7: Activity of Orai1 double/triple point mutants always containing the K85E mutation.** a) – c) Block diagram of current densities of Orai1 K85E L130S, Orai1 K85E H134A, Orai1 K85E F136S, Orai1 K85E L185A F250A, Orai1 K85E V181K, Orai1 K85E V181A, Orai1 K85E P245L and Orai1 K85E S239C compared to the corresponding constitutively active Orai1 mutants (in the absence of K85E). d) Intensity of plasma membrane fluorescence of selected YFP-labelled Orai1 mutants in a) – c). e) – h) Intensity plots of STIM1-OASF co-expressed with Orai1 H134A compared to Orai1 K85E H134A (e); Orai1 L185A F250A compared to Orai1 K85E L185A F250A (f); Orai1 V181K compared to Orai1 K85E V181K (g); Orai1 P245L compared to Orai1 K85E P245L(h). i) Block diagram of maximum currents of Orai1-SS-GFP compared to Orai1 K85E-SS-GFP and Orai1 K85E H134A-GFP.

**Supplementary Figure 8: Properties of Orai1 gain- and loss-of-function mutants in the MTR section.** a) Intensity of plasma membrane fluorescence of YFP-labelled Orai1 H134A, S239C, L174D, L174D S239C and Orai1 H134A L174D. b) - e) Intensity plots of STIM1-OASF co-expressed with Orai1 V181K compared to Orai1 E149K V181K (b); Orai1 P245L compared to Orai1 E149K P245L (c); Orai1 L174D P245L and Orai1 S179F P245L (d) and Orai1 S239C compared to Orai1 L174D S239C (e). f) Intensity of plasma membrane fluorescence of YFP-labelled Orai1 V181K, Orai1 E149K V181K, Orai1 P245L and Orai1 E149K P245L, Orai1 L174D P245L and Orai1 S179F P245L. g) – i) Intensity plots of STIM1-OASF co-expressed with Orai1 H134A compared to Orai1 H134A S179F and Orai1 H134A L174D (g); Orai1 F136S compared to Orai1 F136S S179F (h) and Orai1 H134A compared to Orai1 H134A L174D (i). j) Intensity of plasma membrane fluorescence of YFP-labelled Orai1 H134A compared to Orai1 H134A L174D, Orai1 H134A S179F and Orai1 F136S compared to Orai1 F136S S179F. k) Block diagram of STIM1 mediated currents of Orai1 H134A compared to Orai1 H134A E149K and Orai1 H134A I148S. l) Intensity plots of STIM1-OASF co-expressed with Orai1 H134A compared to Orai1 H134A E149K and Orai1 H134A I148S. m) Intensity of plasma membrane fluorescence of YFP-labelled Orai1 H134A compared to Orai1 H134A I148S, Orai1 H134A E149K and Orai1 L185A F250A compared to Orai1 L185A F250A L174D. n) Block diagram of STIM1 mediated current densities of Orai1 L185A F250A compared to Orai1 L174D L185A F250A. o) Intensity plots of STIM1-OASF co-expressed with Orai1 L185A F250A compared to Orai1 L174D L185A F250A.

**Supplementary Figure 9: The effect of LoF mutants on hydration profile of Ora1 H134A.** a-f) Pore hydration profile for wild type Orai1, Orai1 H134A and Orai1 H134A double mutants. The number of water molecules is given as a function of the distance from the selectivity filter. Hydration profiles are given for wild type in dotted black line where the gray shaded areas correspond to the standard deviation of the mean. Solid black, red, purple, blue orange and green lines correspond to Orai1 wild-type, Orai1 H134A, Orai1 K85E H134A, Orai1 H134A E149D, Orai1 H134A L174D and Orai1 H134A S239W. The profiles were calculated t = 200 – 250 ns, 250 – 300 ns, 300 – 350 ns and 350 – 400 ns. Positions of the carbon α of the residues delineating the pore are given on the top axis with distance from the selectivity filter shown on the bottom.

**Supplementary Figure 10: Orai1 dimers containing one wild-type and one mutant subunit with H134A or P245L form mainly store-operated active channels. a) – b)** Time course and block diagrams of Orai1 dimer whole cell current densities in the absence compared to the presence of STIM1. Orai1 dimer mutants represents Orai1 – Orai1 H134A (a), Orai1 – Orai1 P245L (b).

**Supplementary Figure 11: Distances of residues in the cytosolic triangle relative to the hydrophobic gate and relative to each other.** a) Top: Distances from the hydrophobic gate for the carbon alpha of different residues (83, 85, 149, 173 respectively in blue, cyan, green and red) projected along the normal of the pore as a function of the simulation time. For each residue, the mean over each subunit was calculated and the shaded area corresponds to the standard deviation. Bottom: Distance from the hydrophobic gate for the charged moieties (carboxyl, amino or guanidinium) of different residues (83, 85, 149, 173 respectively in blue, cyan, green and red) projected along the normal of the pore as a function of the simulation time. For each residue, the mean over each subunit was calculated and the shaded area corresponds to the standard deviation. b) Distances between charged groups of residues 83, 85, 149, 173 for wild-type and some of its mutants as a function of the simulation time. Distances between 83-173, 83-149, 85-173 and 85-149 are plotted in blue, cyan, green and red, respectively. For each distance, the mean over each subunit was calculated and the shaded area corresponds to the standard deviation. The coloring of the borders of each graph indicate which hOrai1 we are looking at. Top left in black, wildtype. Top right in cyan, E149K. Bottom left in pink, K85E. Bottom right in orange, E173K. c) (left) Side view of an Orai subunit showing the interactions with neighboring TM domains within the (solid) and of its adjacent subunit (ghost). TM1, TM2 and TM3 are represented in glassy ribbon material in red, cyan and green respectively. Residues K85, R83, E149 and E173 are colored in blue, cyan, orange and red, respectively. (right) Top view of the entire Orai1 channel complex with K85, R83, E149 and E173 highlighted with glassy ribbon material in blue, cyan, orange and red.

**Supplementary Figure 12: Pore profile of Orai1 point mutants and intensity profiles of STIM1 C-terminal fragment when co-expressed with Orai1 mutants.** a – c) Molecular dynamics simulations demonstrate the changes of the pore geometry in the presence of mutations within the cytosolic triangle. a) Superposition of snapshots at *t* = 200-250 ns from MD simulations of WT (gray) and K85E (purple) (a), E149K (cyan) (b), L81K K85E (yellow) (c) as viewed from the top. Respective pairs of diagonal subunits viewed from the side (bottom). d -f) Intensity plots of STIM1-OASF or STIM1 OASF-L251S co-expressed Orai1 K85E in comparison to Orai1 L81K K85E (d), Orai1 K85E E173K (e) and Orai1 R83E E149K (f).

**Supplementary Figure 13: Maximum current densities of single and multiple point mutants containing substitutions of the residues in the cytosolic triangle.** a) Block diagram showing STIM1 mediated maximum currents of Orai1 L81K K85E, S82K K85E, K85E A88K and K85E S89K. b) Block diagram showing maximum current densities of Orai1 mutants containing substitutions of the residues in the cytosolic triangle. Scheme with the residues of the cytosolic triangle highlighting attracting and repulsing forces. Right to the schemes the number of attracting and repulsing forces is indicated. As long as 2 repulsing forces are present the respective mutant shows STIM1 mediated store-operated activation.

**Supplementary Figure 14: Role of L174 in the CETR**. a) *(left)* Superposition of snapshots at *t* = 200-250 ns from MD simulations of WT (gray) and L174D (yellow) as viewed from the top. *(middle)* Pairs of diagonal subunits viewed from the side. *(right)* Pore hydration profile of wild type Orai1 and Orai1 L174D. The number of water molecules is given as a function of the distance from the selectivity filter. Hydration profiles are given for wild type in dotted black line where the gray shaded areas correspond to the standard deviation of the mean. Scattered and solid orange lines correspond to Orai1 L174D at t = 150 - 200 ns and t = 200 – 250 ns, respectively. Positions of the carbon α of the residues delineating the pore are given on the top axis with distance from the selectivity filter shown on the bottom. b) Top and bottom: Salt-bridges between residues at position 173 (top) or position 149 (bottom) and K85 either in wild-type Orai1 or Orai1 H134A. c) Time course block diagram of STIM1 mediated current densities of Orai1 E173K, Orai1 L261D and E173K L261D. Block diagram shows STIM1 mediated maximum current densities of Orai1 E173K, Orai1 L261D, Orai1 E173K L261D, Orai1 K85E E173K L261D, Orai1 L174D L261K and Orai1 L174K L261D.

**Supplementary Figure 15: Operator symbols and cytosolic triangles in Orai1.** a) Schematic representation of a logical input AND and OR gates with two inputs (A, B) and one output (C) and corresponding truth tables. All inputs and the outputs can only take the values 0 or 1. AND gate: Only as long as both inputs are 1 the output is 1. If one of the two inputs is 0, the output is 0. OR gate: If either one of the two inputs is 1, the output is 1. Only if both inputs are 0, the output is 0. b) Description of the contribution of the cytosolic triangles to the Orai1 activation mechanism by the use of the operator symbols for the AND and OR gates. Always two salt-bridge interactions, either, R83-E149 and K85-E173, K85-E173 and K85-E149 or R83-E149 and K85-E149, can be viewed as an individual AND gate. One of the two salt-bridge pair, thus one AND gate, is required to allow pore opening. Hence, the three AND gates can be combined by an OR gate. These requirements are also described in the truth table in (b).

### Supplementary Figure 1

**Screen across residues in TM2, TM3 and TM4 located in close-proximity**

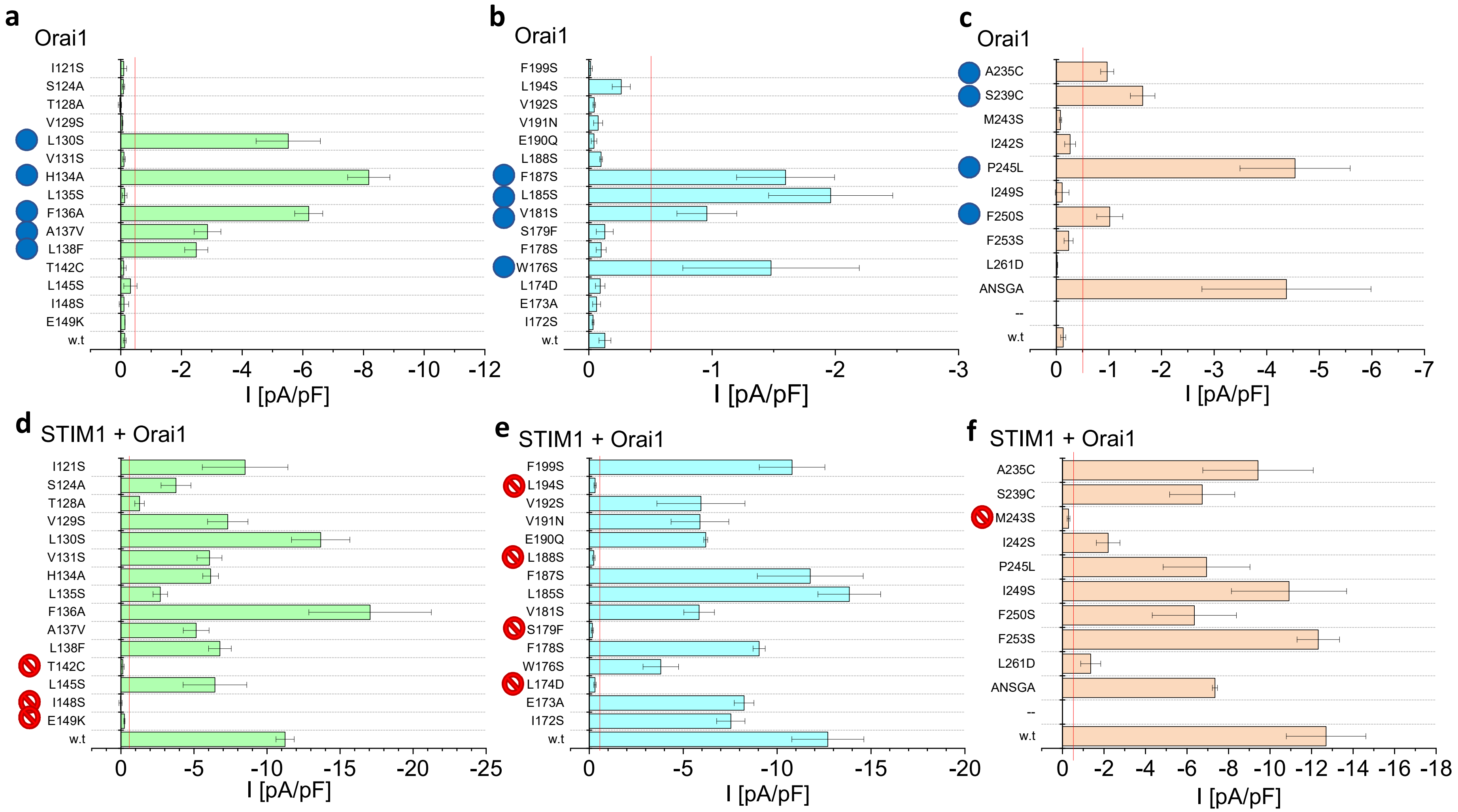

### Supplementary Figure 2I

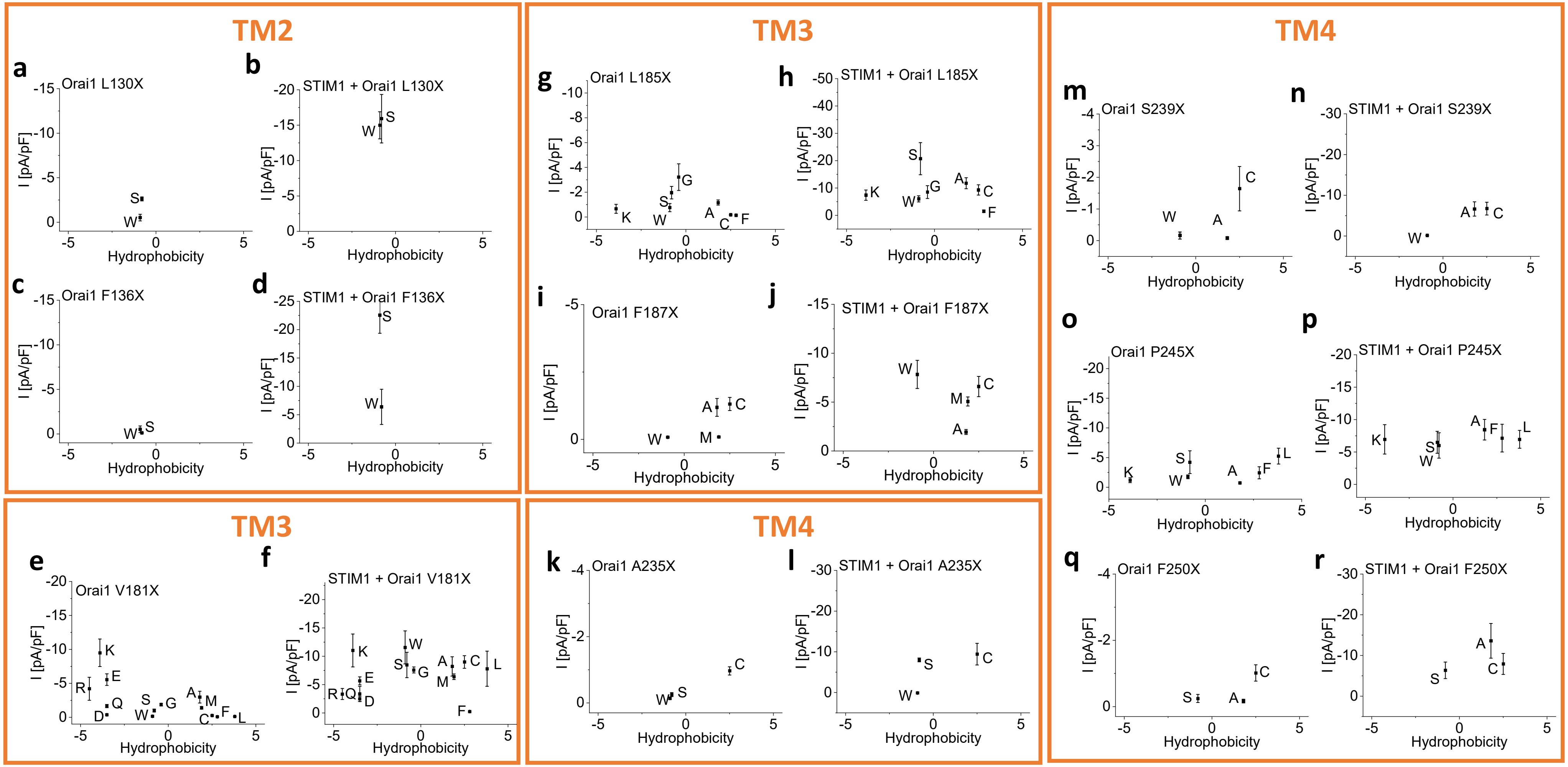

### Supplementary Figure 2II

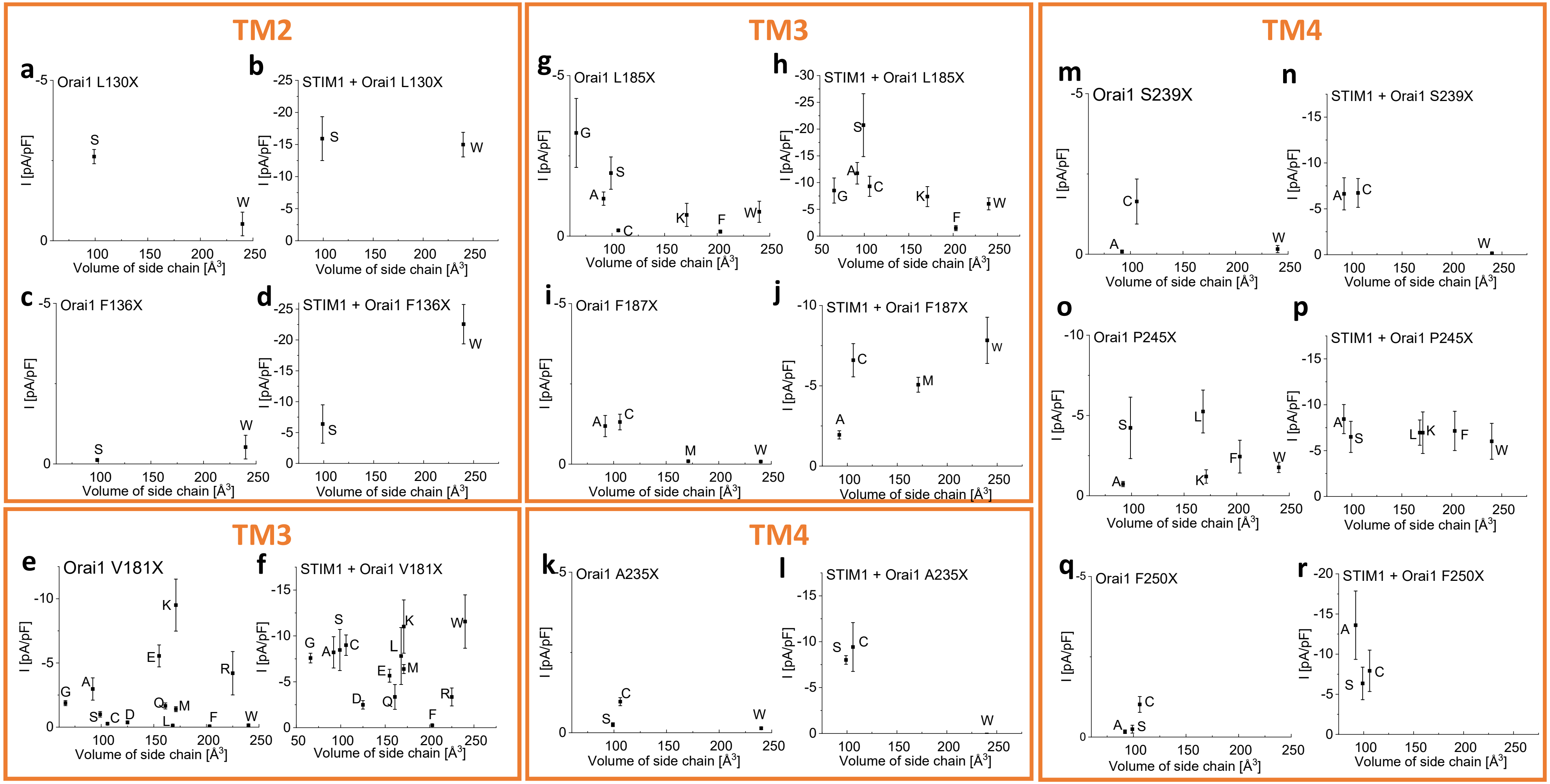

### Supplementary Figure 3

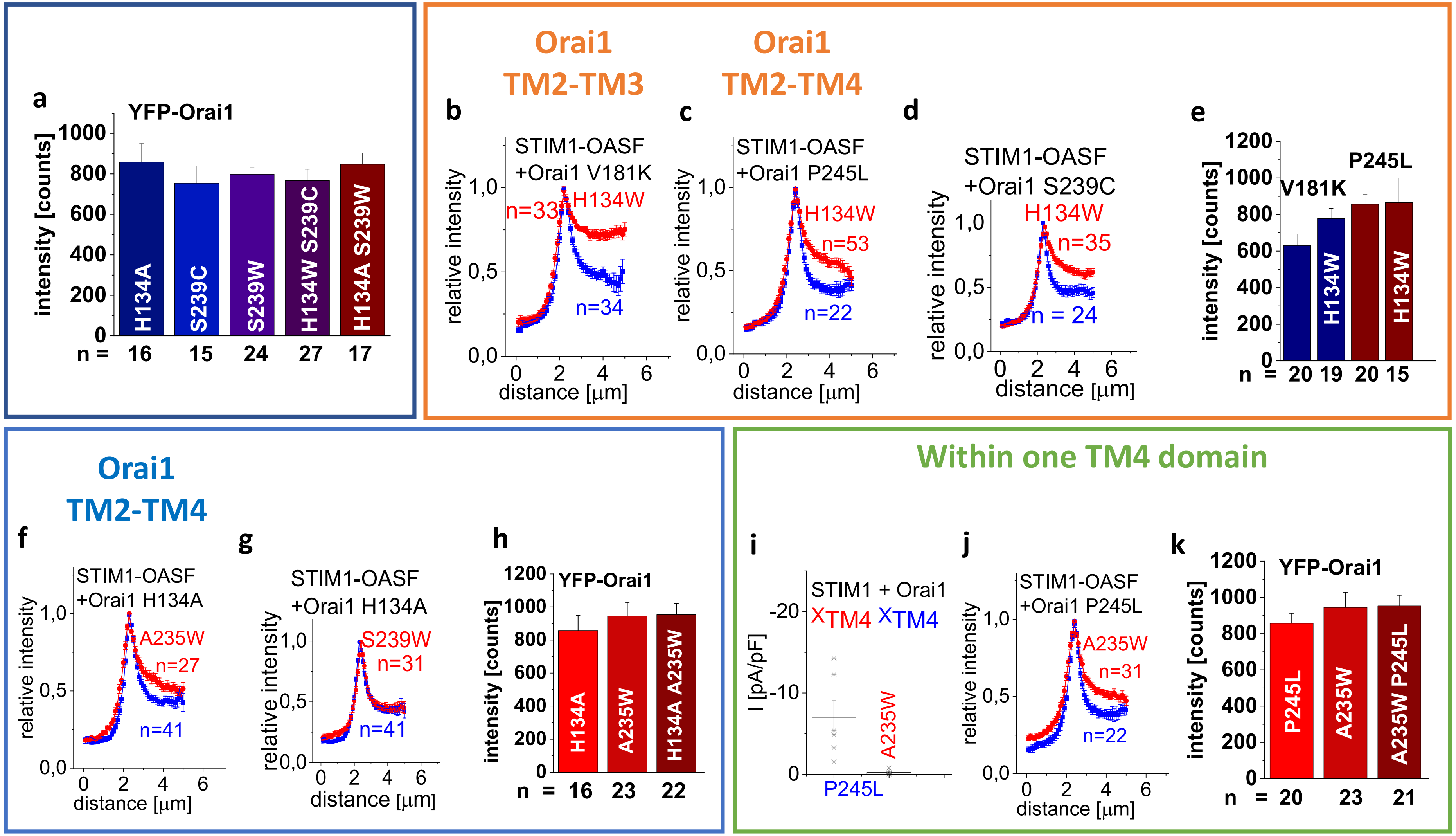

### Supplementary Figure 4

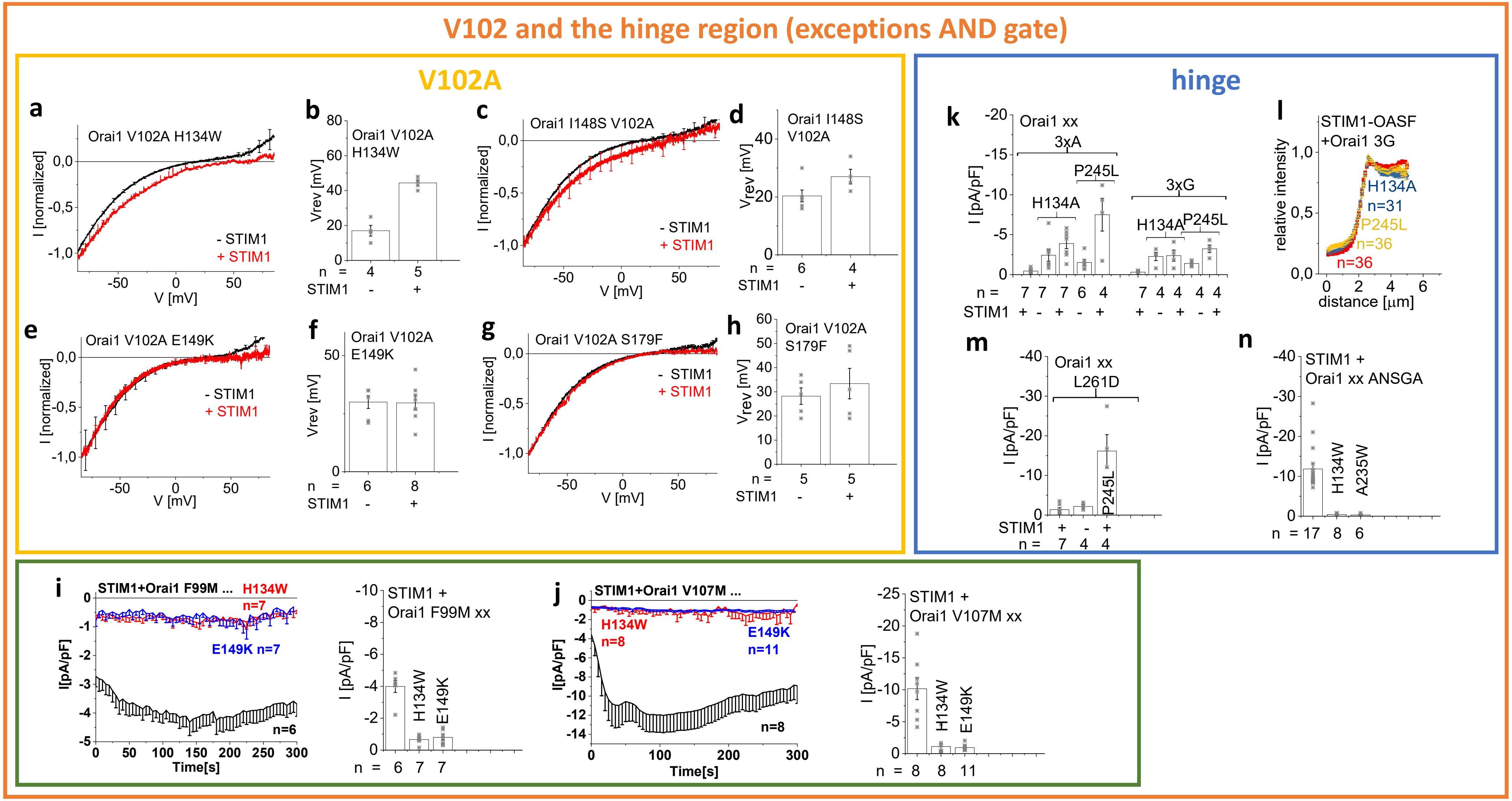

### Supplementary Figure 5

Additional novel **loss-of-function mutants** in the MTR

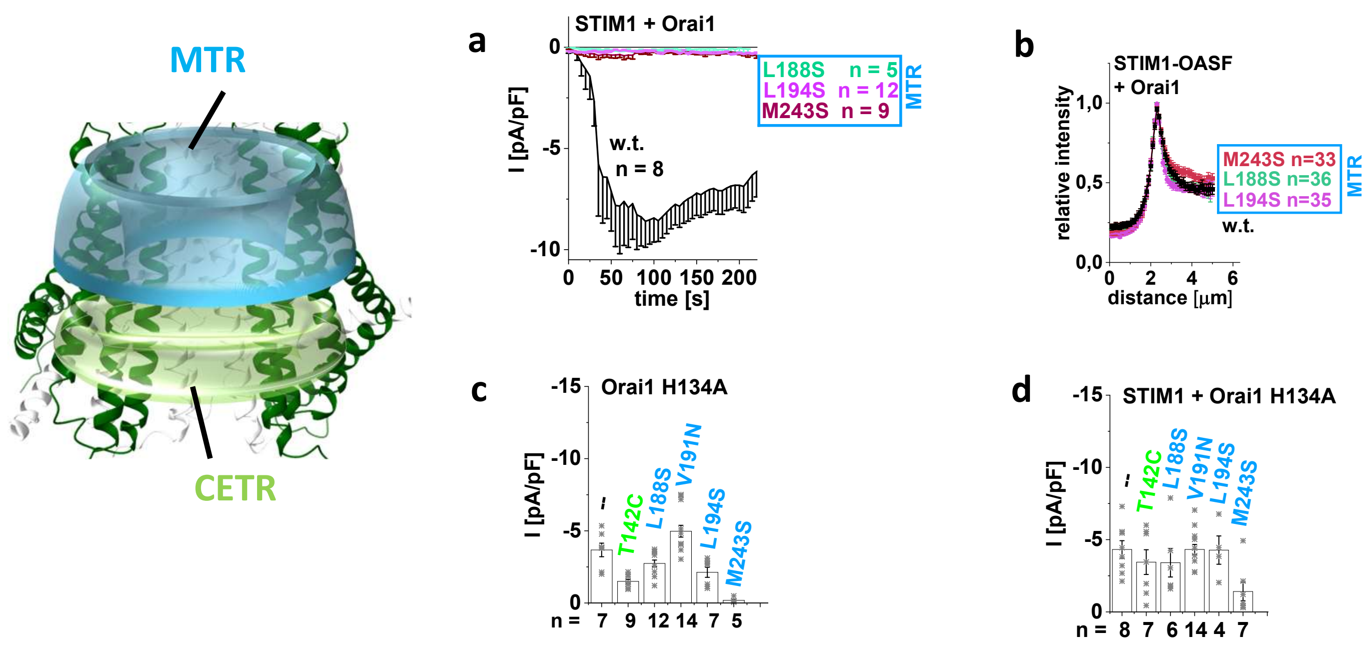

### Supplementary Figure 6

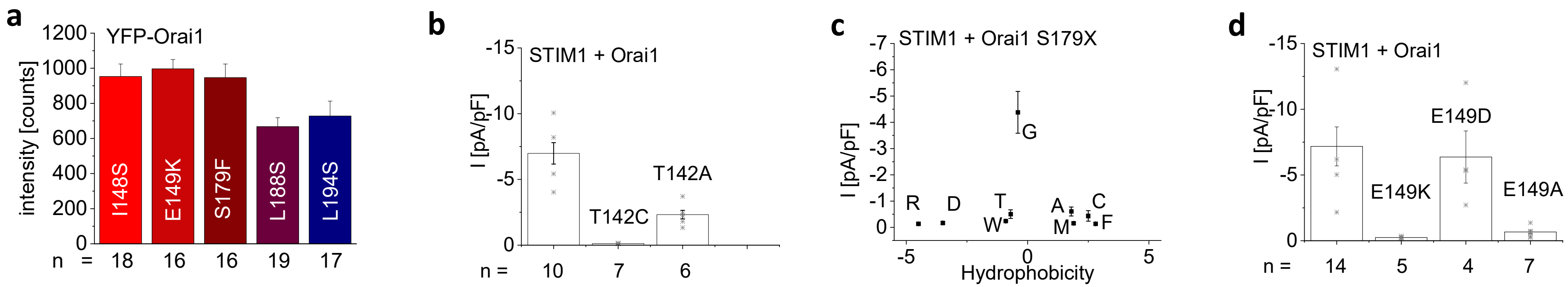

### Supplementary Figure 7

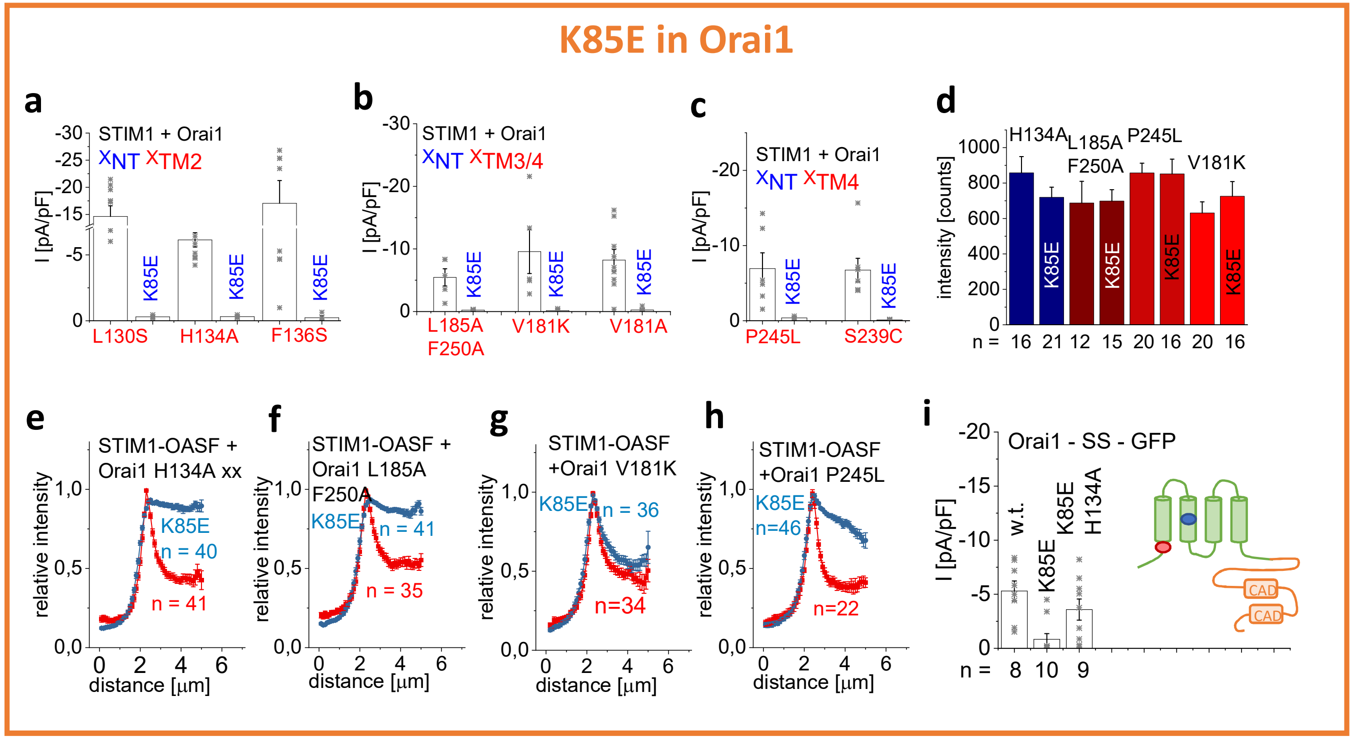

### Supplementary Figure 8

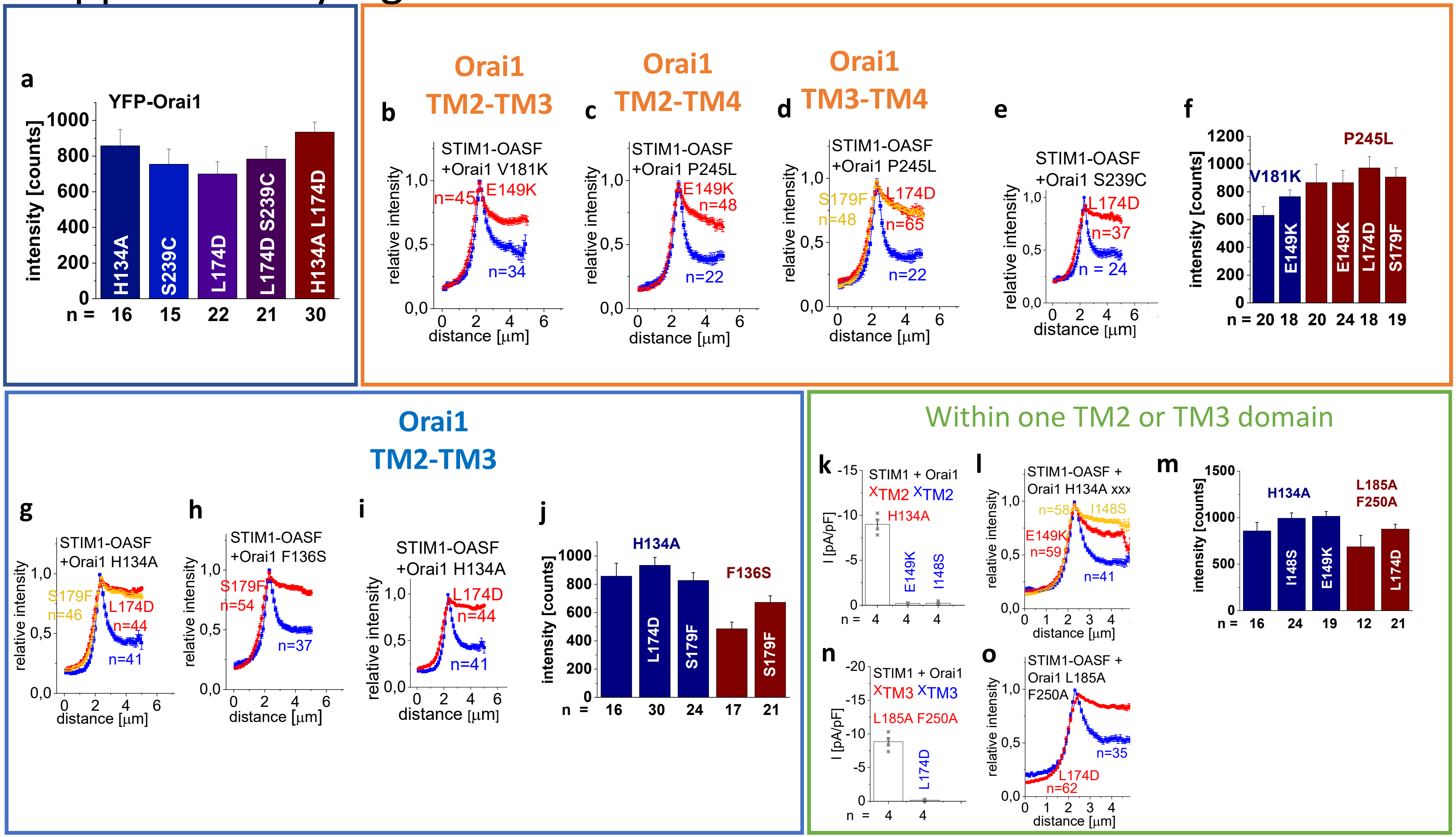

Supplementary Figure 9

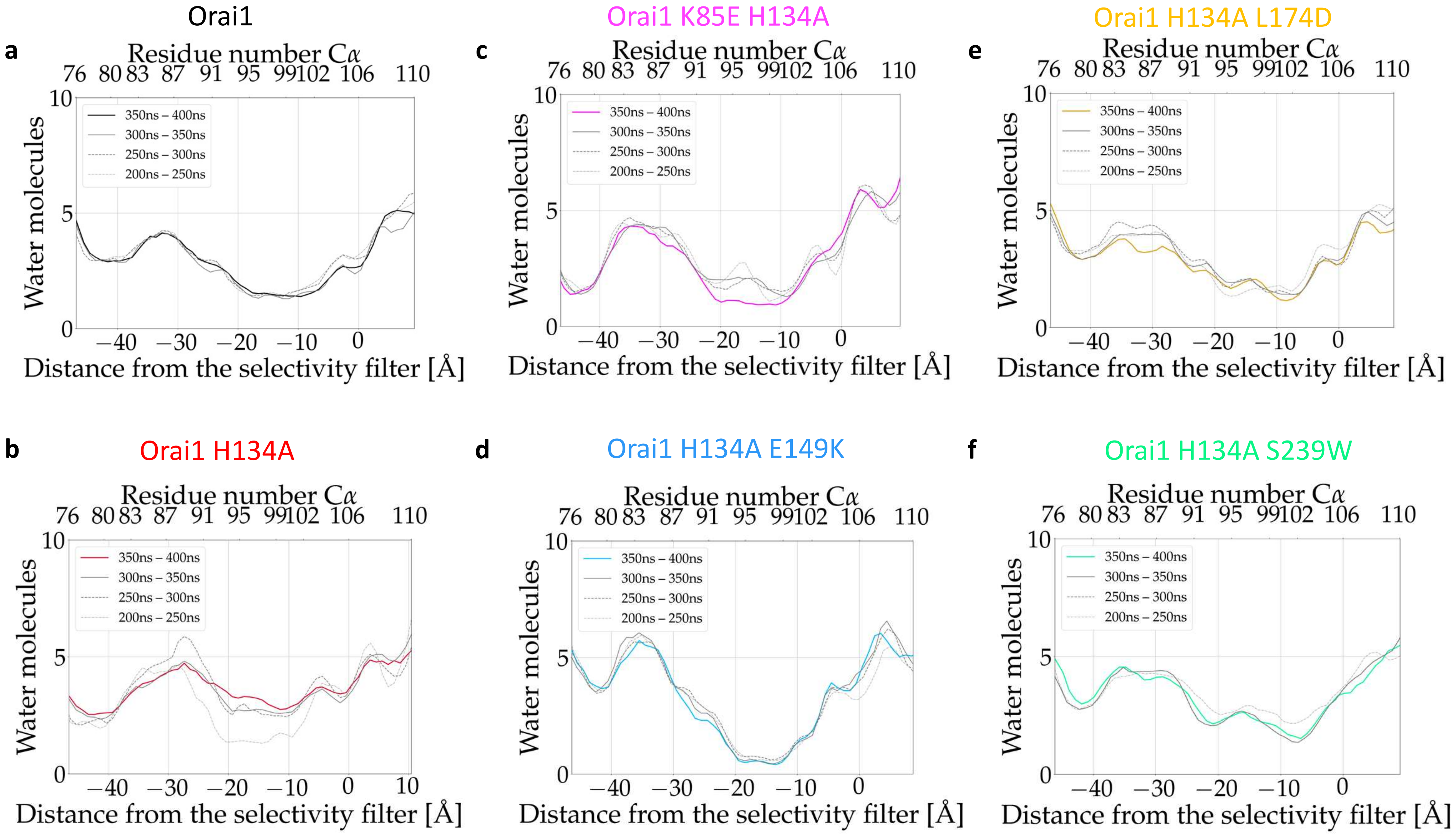

### Supplementary Figure 10

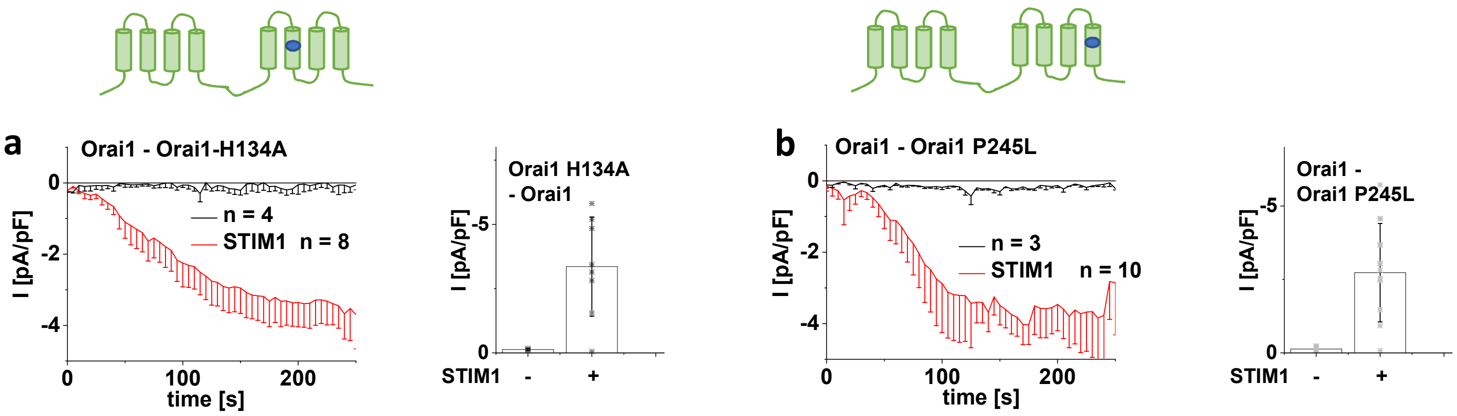

### Supplementary Figure 11

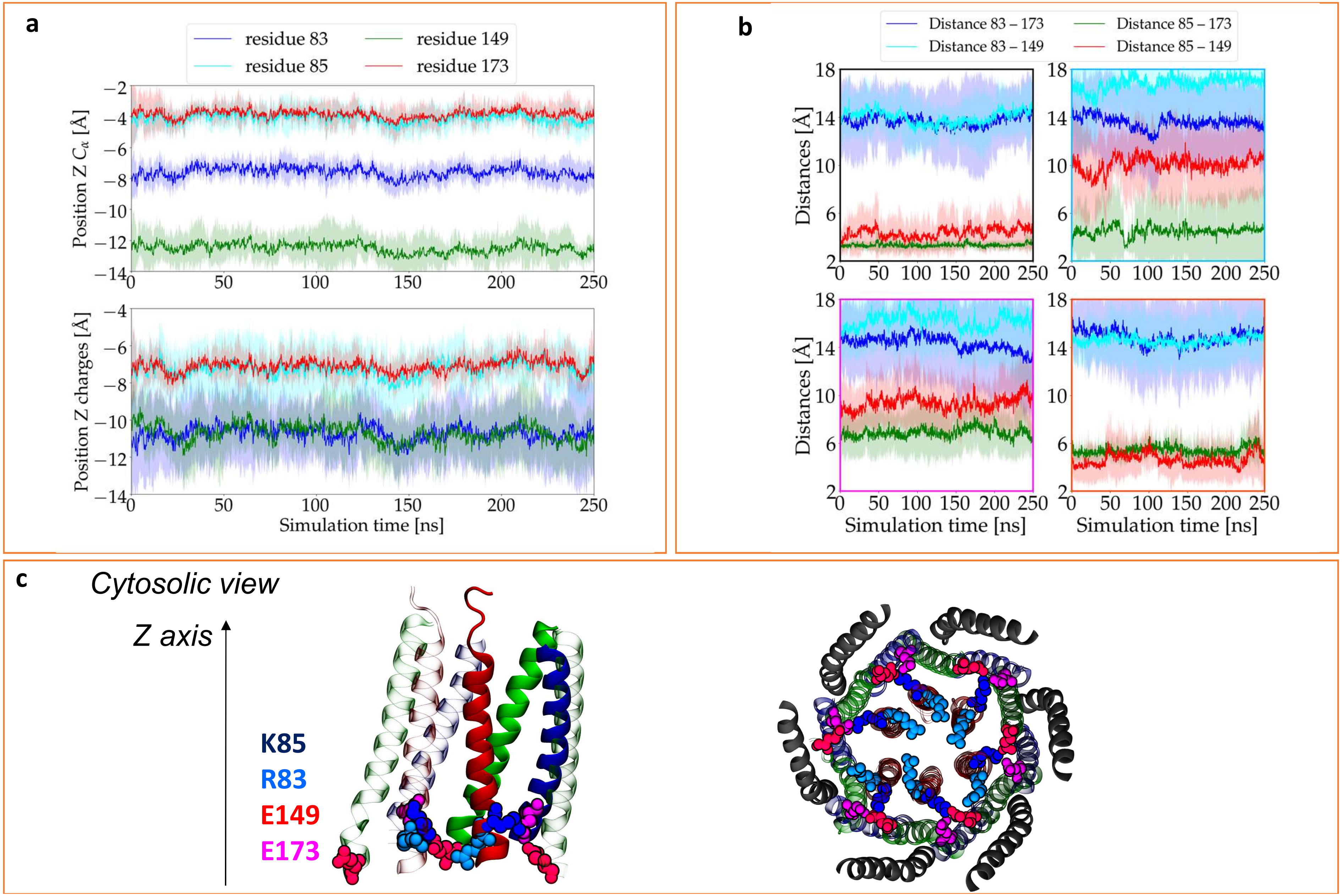

Supplementary Figure 12

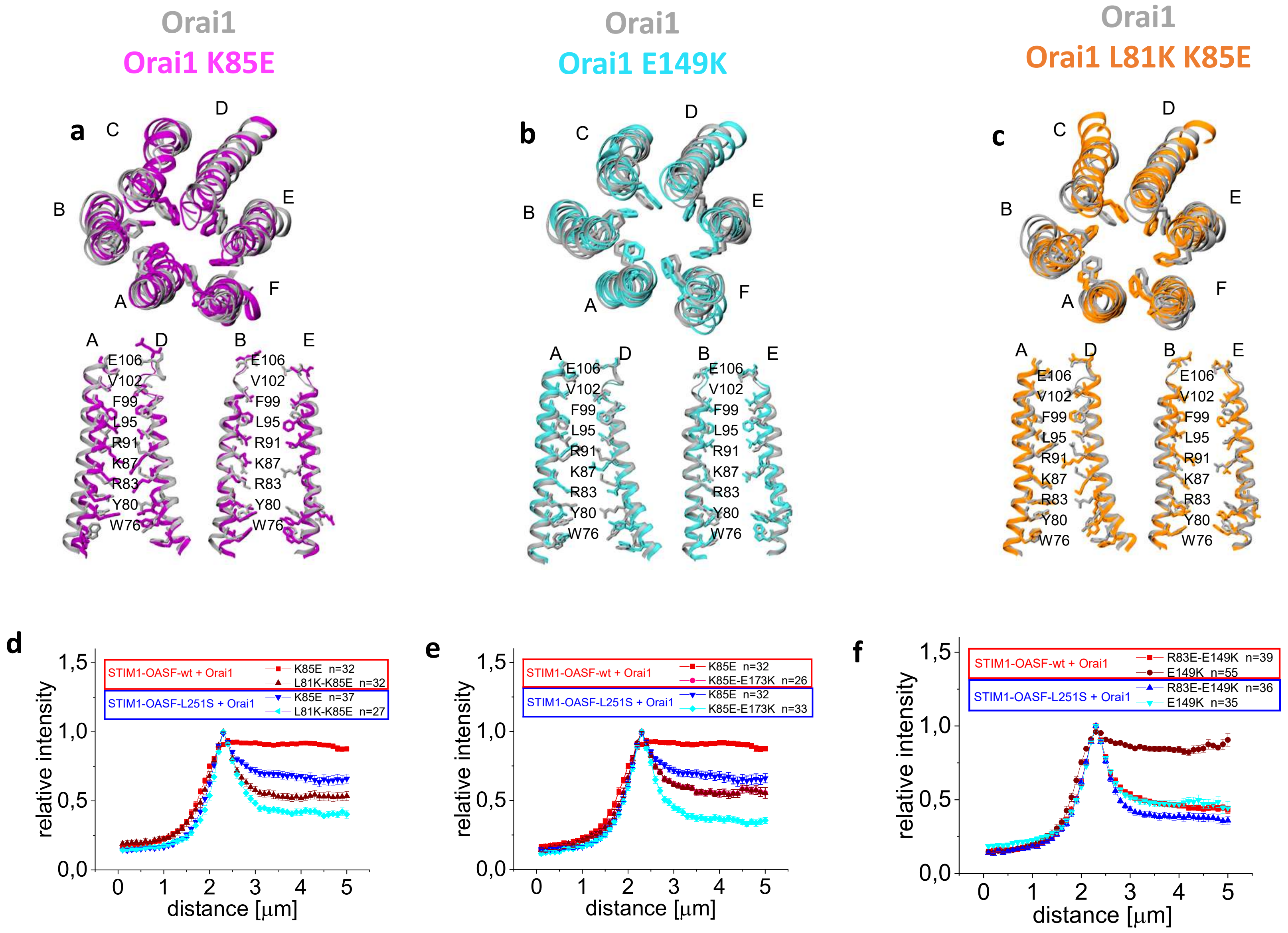

### Supplementary Figure 13

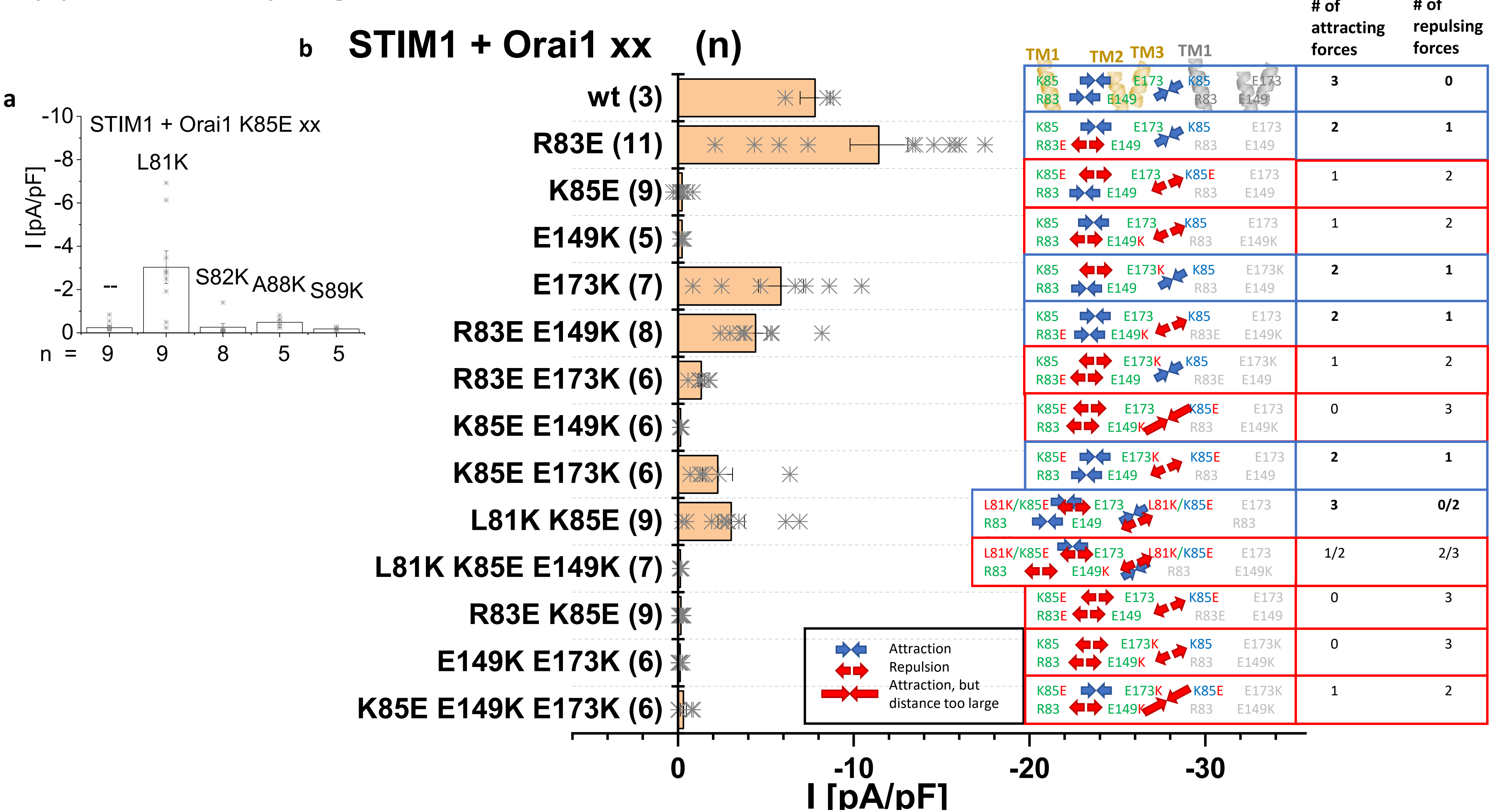

### Supplementary Figure 14

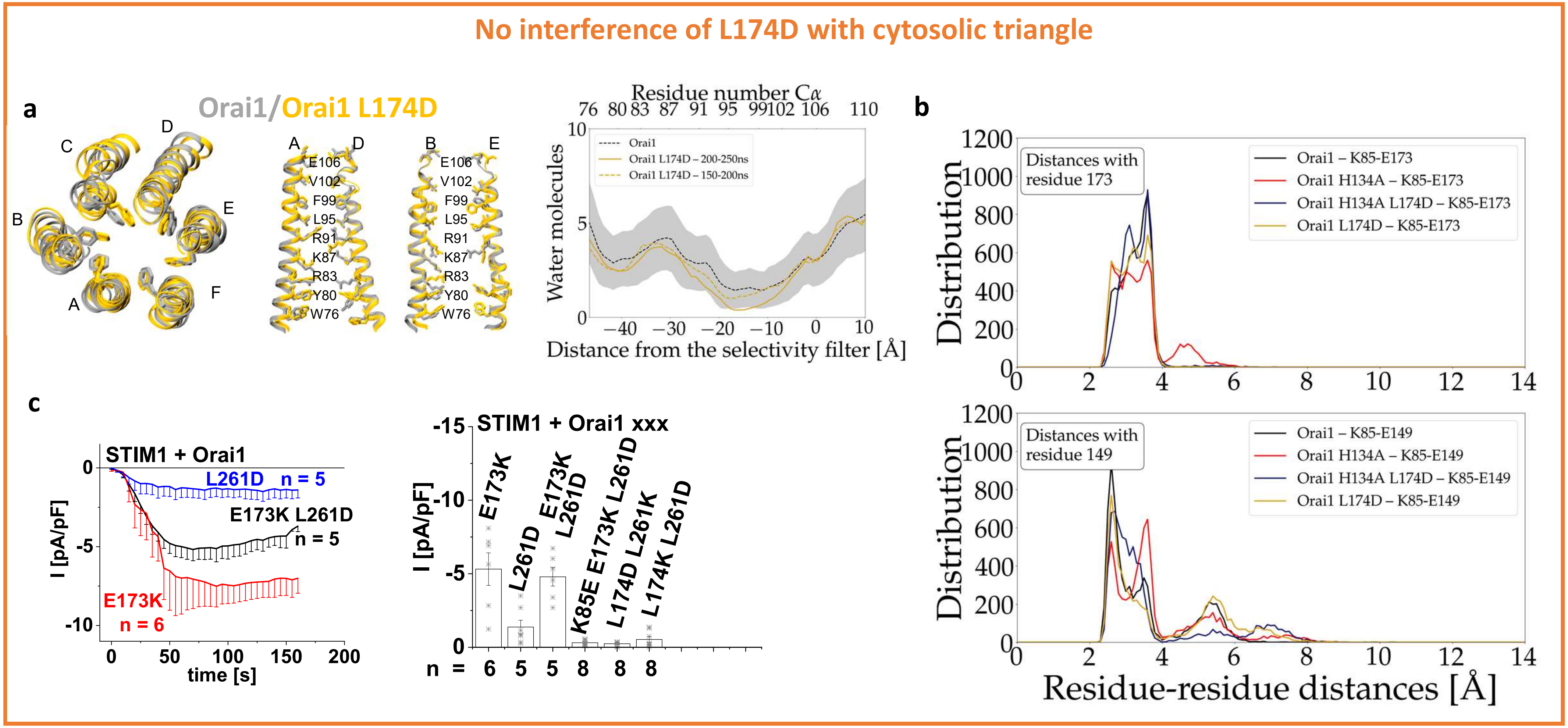

Supplementary Figure 15

**a**

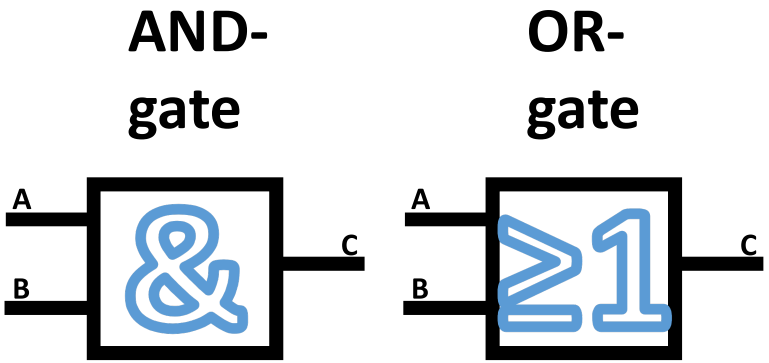


| A | B | C |
|---|---|---|
| 0 | 0 | 0 |
| 1 | 0 | 0 |
| 0 | 1 | 0 |
| 1 | 1 | 1 |

| A | B | C |
|---|---|---|
| 0 | 0 | 0 |
| 1 | 0 | 1 |
| 0 | 1 | 1 |
| 1 | 1 | 1 |

**b** **At least two salt-bridge bonds in the CTS guarantee Orai function**

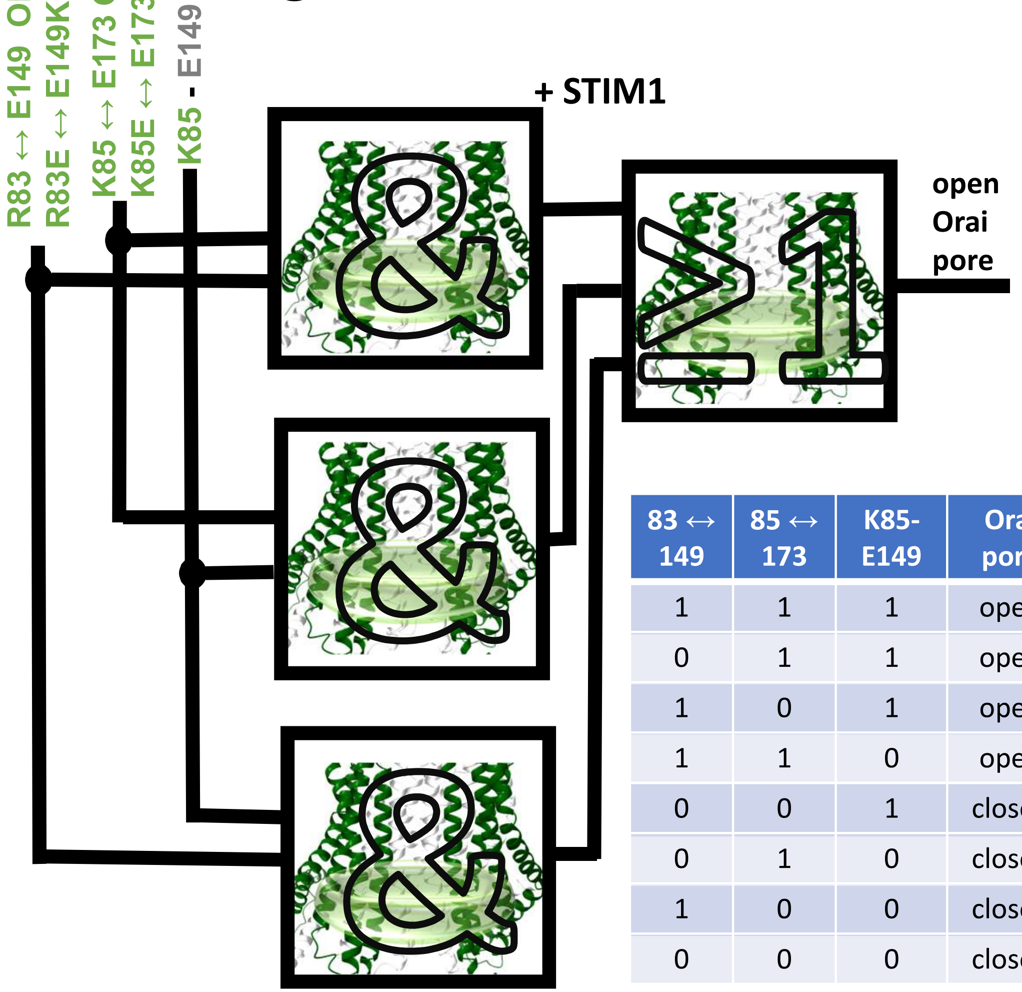


| 83 ↔ 149 | 85 ↔ 173 | K85-E149 | Orai pore |
|---|---|---|---|
| 1 | 1 | 1 | open |
| 0 | 1 | 1 | open |
| 1 | 0 | 1 | open |
| 1 | 1 | 0 | open |
| 0 | 0 | 1 | closed |
| 0 | 1 | 0 | closed |
| 1 | 0 | 0 | closed |
| 0 | 0 | 0 | closed |
